# Genetically encoded Boolean logic operators to sense and integrate phenylpropanoid metabolite levels in plants

**DOI:** 10.1101/2024.01.05.574427

**Authors:** Savio S. Ferreira, Mauricio S. Antunes

## Abstract

- Synthetic biology has the potential to revolutionize biotechnology, public health and agriculture. Recent studies have shown the enormous potential of plants as chassis for synthetic biology applications. However, tools to precisely manipulate metabolic pathways for bioproduction in plants are still needed.
- We have adapted bacterial allosteric transcription factors (aTFs) to control gene expression in plants in a ligand-specific manner. The aTFs used here function as transcription repressors of semi-synthetic promoters, and aTF activity is regulated by specific plant metabolites, especially phenylpropanoid-related molecules. Using these aTFs, we also designed synthetic genetic circuits capable of computing Boolean logic operations.
- Three aTFs, CouR, FapR and TtgR, were able to achieve ∼95% repression of their respective target promoters. For TtgR, a 6-fold de-repression could be triggered by inducing its ligand (naringenin) accumulation, showing its use as biosensor. Moreover, we designed synthetic genetic circuits that use AND, NAND, IMPLY and NIMPLY Boolean logic operations and integrate metabolite levels as input to the circuit.
- We showed that biosensors can be implemented in plants to detect phenylpropanoid-related metabolites and activate a genetic circuit that follows a pre-defined logic, demonstrating their potential as tools for exerting control over plant metabolic pathways and facilitating the bioproduction of natural products.

## Introduction

Synthetic biology is a rapidly evolving field at the intersection of various disciplines, harnessing principles from engineering, genetics, molecular biology, and others, to create innovative solutions for agriculture, public health, biotechnology, and sustainability (Kitada *et al.,* 2018; Scown & Keasling 2022; Yang & Reyna-Llorens 2023). Several synthetic biology-derived products are now commercially available to benefit our daily lives, with many more to come in the near future (Voigt 2020; Meng & Ellis 2020). Most of the research in synthetic biology is currently carried out in microbial systems, as they have a wide availability of biotechnological tools, smaller genomes and shorter life cycles (Burnett & Burnett 2020; Liu *et al.,* 2023). On the other hand, synthetic biology in plants still lags behind due to inherent properties such as longer life cycles, multicellularity and diversity of cell types, as well as a lower availability of biotechnological tools and standard genetic parts (Burnett & Burnett 2020; Liu *et al.,* 2023). Despite these challenges, recent plant synthetic biology studies have made substantial progress (Patron *et al.,* 2015; Khakhar *et al.,* 2018; Bernabé-Orts *et al.,* 2020; Brophy *et al.,* 2022; Lloyd *et al.,* 2022; Selma *et al.,* 2022; Anderson *et al.,* 2023; Ranawaka *et al.,* 2023), as plants are widely recognized as an important chassis for engineering efforts towards a sustainable bioeconomy.

Plant synthetic biology seeks to not only improve crop yield by manipulating physiological aspects to allow more efficient use of resources and better resilience toward stresses, but also to engineer plants to produce high-value chemicals and even to introduce entirely novel functions, such as sensing (Patron 2020; Yang *et al.,* 2022; Yang & Reyna-Llorens 2023). Among these aspects, much attention has been given to manipulating plant specialized metabolism, as plants naturally produce an astonishing array of unique chemicals with potential for biotechnological applications (Garagounis *et al.,* 2021; Liu *et al.,* 2023). The phenylpropanoid pathway yields a variety of phenolic compounds, such as flavonoids and stilbenes, with diverse applications, including antimicrobial and antioxidant properties, food additives and fragrances (Neelam *et al.,* 2020). Notably, significant effort has also been devoted to transposing plant specialized pathways to microbial systems to produce these metabolites by fermentation (Cravens *et al.,* 2019; Sun *et al.,* 2022). Nonetheless, reconstructing an entire metabolic pathway in a heterologous system is not a straightforward task, as it may require dozens of genes to be stacked. Furthermore, protein misfolding, metabolite compartmentalization requirements, lack of plant-specific co-factors and precursors and antimicrobial activity of the plant metabolite may result in failure of the introduced pathway in the microbial host (Lin *et al.,* 2019; Keita *et al.,* 2022). On the other hand, plants have several features that make them ideal chassis for bio-production of specialized metabolites, including large biomass accumulation, lack of reliance on carbon sources, compartmentalization (organelles, tissues, cell types), shared pathways, transient gene expression systems, and others (Molina-Hidalgo *et al.,* 2021; Liu *et al.,* 2023).

As we move towards engineering plants as biofactories, synthetic biology tools will be required to control and optimize plant metabolic pathways to unlock the promise of sustainable bioproduction. Given this scenario, synthetic genetic circuits that are capable of sensing signals or stimuli and integrating this information dynamically are key to obtaining precise modulation of a physiological function or metabolic pathway (Bull & Khakhar 2023). Genetically encoded Boolean logic gates are devices that can be deployed to process multiple inputs (signals, molecules, stimuli) and implement digital-like computations, endowing plants with decision-making abilities. In Boolean logic, the values of the variables (inputs, outputs) are either true or false (represented numerically by 1 or 0, respectively). These values can also be used to represent the presence (1) or absence (0) of a molecule or a signal. Input values are computed following Boolean operations such as conjunction (AND), disjunction (OR), and negation (NOT), resulting in an output. Based on the applied logic, the output is true only when specific combinations of inputs are met, and the overall operation is represented by a truth table. Logic gates are the simplest form of information processing, based on the presence or absence of signals, and many natural genetic networks in living cells can be modeled using Boolean logic (Ferreira *et al.,* 2023; Koukara & Papadopoulou 2023). Simple logic gates typically integrate two inputs; however, multiple inputs can be computed by layering logic operations, increasing processing power and reliability of the system to respond according to user-defined rules (Koukara & Papadopoulou 2023; Ferreira *et al.,* 2023).

In the context of controlling metabolic pathways, one effective strategy involves equipping genetic circuits with the capability to sense specific molecules, which can then serve as inputs for Boolean logic operations, allowing for precise modulation and regulation of cellular processes. Allosteric transcription factors (aTFs) are sensor proteins commonly found in bacteria whose transcriptional activity is modulated by a specific metabolite (ligand or effector) or closely related metabolites (Li *et al.,* 2023). These aTFs usually have two well-defined domains, a DNA binding domain (DBD) and a ligand binding domain (LBD), and belong to different protein families such as MarR, AraC, LacI, TetR, GtnR, FapR, LysR, among others (d’Oelsnitz *et al.,* 2022; Li *et al.,* 2023) with no significant sequence identity across families. In general, aTFs bind to a specific sequence (response element, RE) in the operator of target genes and their affinity for the RE is controlled by the ligand (Fig. **1a**). aTFs that function as transcriptional repressors bind to the RE in the absence of the ligand, repressing expression of downstream genes. Upon ligand binding, the aTF undergoes conformational changes causing its release from the RE, thus allowing transcription to occur (Fig. **1a**). Several natural aTFs have been characterized as responsive to plant metabolites, as they usually regulate genes responsible for the degradation of these plant-derived chemicals to be used as carbon sources or to detoxify the cell (Alguel *et al.,* 2007; Otani *et al.,* 2016). Among these, aTFs like TtgR, CouR, PadR, FdeR and FerC are responsive to phenylpropanoid-related molecules (Ferreira & Antunes 2021), making them good sensor candidates to engineer plants with the goal of manipulating the synthesis of phenylpropanoid-derived metabolites. In fact, several of these sensors have been deployed in microbial systems to engineer phenylpropanoid producing strains (Zhang *et al.,* 2023). These aTFs can sense the metabolic status of the microbial cell, thereby providing dynamic regulation of an introduced pathway based on metabolite levels. This approach allows better control of the carbon flux towards the pathway and therefore avoids metabolic burden, resource competition and toxicity, leading to significant yield increases (Xu *et al.,* 2014; Zhou *et al.,* 2021).

**Fig. 1.**
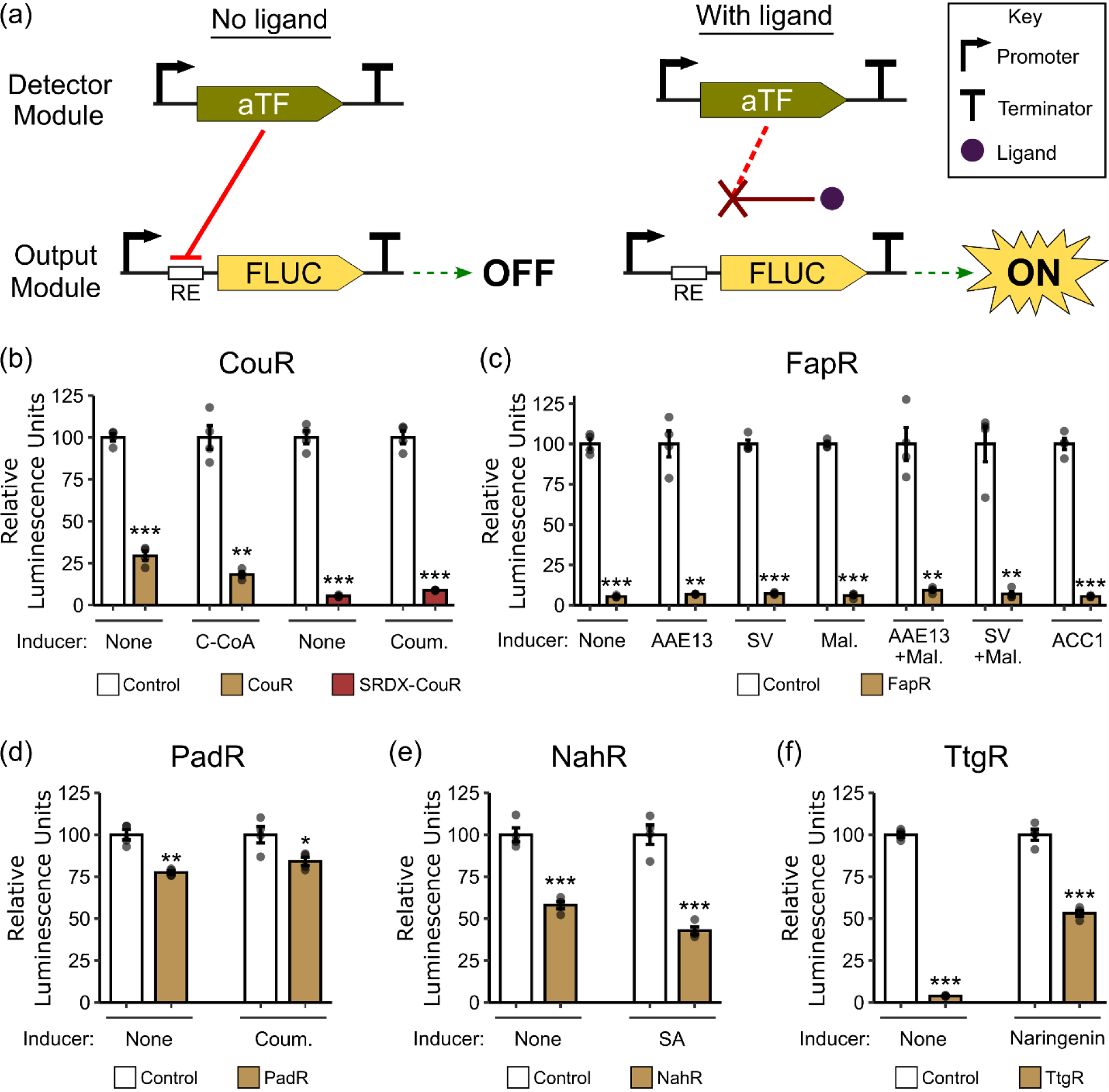
Biosensor systems based on repressor aTFs and transient assays in Arabidopsis protoplasts. (a) Schematic representation of the biosensor devices used in this work. The detector module constitutively expresses the aTF. In the absence of the ligand (left), the aTF binds to the response element (RE) in the target promoter of the reporter gene, Firefly Luciferase (FLUC), in the output module, repressing its expression. When the ligand is available (purple circle), it bids to the aTF hindering the interaction with the RE, thus allowing FLUC expression. (b-f) Transient assays in Arabidopsis protoplasts testing the ability of each aTF, CouR (b), FapR (c), PadR (d), NahR (e) and TtgR (f), to repress a CaMV 35S promoter harboring their respective RE and to de-repress the system by adding specific inducers. Inducers include enzymes responsible for the biosynthesis of the ligands, precursors of the ligands, or the ligands themselves. Chemical inducers were added to a final concentration of 10 µM. Relative Luminescence Units are the ratio of Firefly (FLUC) and Renilla (REN) Luciferases, normalized by their respective control of each experiment. Error bars = SEM, n = 4, dots represent individual replicates. Statistical significance was determined by 2-tailed Student T-test comparing control to the respective repressed sample (\**p*<0.05, \*\**p*<0.01 \*\*\**p*<0.001). Abbreviations: C-CoA, ρ-coumaroyl-CoA; Coum., ρ-coumaric acid; AAE13, Acyl Activating Enzyme13 (malonyl-CoA synthetase); SV, splice variant of AAE13; Mal., sodium malonate; ACC1, acetyl-CoA carboxylase1; SA, salicylic acid.

To develop genetic circuits in plants that can detect specific metabolite levels in real-time and integrate them into a coordinated response, we describe here ligand-dependent biosensors that can repress gene expression. A biosensor system based on TtgR achieved ∼95% repression of a semi-synthetic promoter, and 6-fold de-repression with its ligand. Moreover, we designed synthetic genetic circuits that use different Boolean logic operations and integrate flavonol levels as circuit inputs. To the best of our knowledge, this is the first report of a biosensor in plants to sense the levels of a class of metabolites in real-time and integrate this information into a synthetic genetic circuit following user-defined rules. This is a significant step toward engineering control of metabolic pathways and achieving customized bioproduction in plants.

## Materials and Methods

### Plasmid Cloning

All plasmids were constructed using the Golden Braid 3.0 (GB3.0) platform (Vazquez-Vilar *et al.,* 2017). Sensor protein coding sequences, CaMV 35S promoters harboring the biosensors REs (Tables **S1** and **S2**) and *CHS* gene (AT5G13930) were synthesized by Integrated DNA Technologies or Twist Bioscience, codon-optimized for *Arabidopsis thaliana*. After cloning individual parts into the pUPD2 domestication vector, transcriptional units (TUs) were built following GB3.0 standard protocols, using domesticated parts (promoters, tomato MYB12 gene Solyc01g079620.2, Firefly Luciferase and Renilla Luciferase reporter genes, and terminators) from the GB2.0 kit purchased from Addgene (https://www.addgene.org/kits/orzaez-goldenbraid2/). The *ACC1* gene (Thlar.0414s0007) was cloned from pennycress (*Thlaspi arvense L.*) in three patches, adding point mutations to remove *Esp*3I and *Bsa*I sites present in the wild-type sequence, which is incompatible with the GB3.0 system. *AAE13* (AT3G16170) and its splice variant were cloned from Arabidopsis Col-0 plants. Total RNA was extracted from pennycress or Arabidopsis seedlings using the RNeasy Plant Mini kit (Qiagen) with in-column DNase treatment, and first strand cDNA was synthesized with One-Taq RT-PCR kit (New England Biolabs), according to manufacturer’s protocols. The fragments from *ACC1* and *AEE13* cDNAs were amplified using Phusion Master mix (Thermo Fisher) with standard cycling parameters and the primers listed in Table **S3**. PCR reactions were purified from gel using the Zymoclean Gel DNA Recovery Kit (Zymo Research). Seventy nanograms of each purified fragment were used for Golden Braid domestication in pUPD2. *ACC1*, *CHS*, *AAE13* and splice variant TUs were built using CaMV 35S promoter and NOS terminator domesticated parts as mentioned above.

### Plant Growth

*Arabidopsis thaliana* Col-0 seeds were surface-sterilized with 10% commercial bleach then sowed in Murashige and Skoog (MS) agar plates, kept at 4°C for two days, and transferred to a Conviron ATC26 growth chamber set to 22 °C, 150 *μ*mol m^−2^ s^−1^ light intensity, short-day (10-h light/14-h dark) cycles, and 60-70% relative humidity. After approximately two weeks, seedlings were transferred to pots containing Sunshine Mix #1 (Sun Gro Horticulture) and grown under the same conditions.

*Nicotiana benthamiana* seeds were surface-sterilized as above then sowed in MS agar plates and directly transferred to LED light racks under long-day conditions (16-h light/8-h dark) at 20-22°C. After one week, seedlings were transferred to pots and grown under the same conditions.

### Protoplast Isolation and Transformation

Protoplasts were isolated and transformed essentially as described in Schaumberg *et al*. (2016). Briefly, 6-10 leaves from ∼10-week-old Arabidopsis Col-0 plants were cut into thin strips with a scalpel blade in W5 solution (154 mM NaCl, 125 mM CaCl_2_, 5 mM KCl, 2 mM MES pH 5.7). Leaves were transferred to enzyme solution (0.4 M mannitol, 20 mM KCl, 20 mM MES, pH 5.7, 1.5% (w/v) Cellulase R-10 (Yakult Honsha), 0.4% (w/v) Macerozyme R-10 (Yakult Honsha), 10 mM CaCl_2_, 1 mg/ml BSA), vacuum was applied for a few seconds followed by incubation at room temperature for 3 h at 40 rpm. Digested leaves were filtered in a 70 μm cell strainer, pelleted by centrifugation at 300 xg for 4 minutes and washed twice with 20 ml of ice-cold W5. Cells were then resuspended in MMg solution (0.4 M mannitol, 15 mM MgCl_2_, 4 mM MES pH 5.7) and the concentration was adjusted to 200,000 protoplasts/ml. For each replicate, 3 µg of plasmid DNA for promoter constructs and 6 µg of biosensor construct (or empty plasmid, for controls) were mixed with 50 µl of protoplasts. Cells were transfected with 1:1 volume (DNA+protoplasts:PEG) of PEG 40% solution (40% w/v PEG4000, 0.2 M mannitol, 100 mM CaCl_2_) for 30 minutes. Reactions were stopped by adding twice the volume of W5 and centrifuging at 300 xg for 4 minutes. Transformed protoplasts were resuspended in 200 µl of WI solution (0.5 M mannitol, 20 mM KCl, 4 mM MES pH 5.7) and plated on 96-well plates for overnight incubation at 22°C in the dark. For experiments with inducers, 5 µl of a 40X working solution of the inducer were pipetted to each well containing 200 µl of transformed protoplasts to achieve the final inducer concentration. To make a 500 mM stock solution, naringenin (TCI), ρ-coumaroyl-CoA (MicroCombiChem GmbH) and ρ-coumaric acid (MP Biomedicals) were dissolved in DMSO, while sodium salicylate (Fisher Chemical) and sodium malonate (Sigma Aldrich) were dissolved in water.

### Agroinfiltration

*Agrobacterium tumefaciens* cultures harboring the desired constructs were grown overnight at 30°C and 230 rpm in 10 ml of liquid LB media containing 30 µg/ml gentamicin, 10 µg/ml rifampicin and 50 µg/ml kanamycin or 100 µg/ml spectinomycin, depending on the backbone binary vector used. Cultures were pelleted at 4,700 xg for 10 minutes, resuspended in Infiltration Solution (10 mM MgCl2, 10 mM MES pH 5.7, 150 µM acetosyringone) to O.D._600_ = 2.0-4.0, and incubated at room temperature for 3-4 hours. Bacterial suspensions were mixed according to each treatment and desired final O.D._600_ and infiltrated into 4-5-week-old *N. benthamiana* plants using a 1-mL needleless syringe. To improve transient expression, the P19 silencing suppressor under control of a 35S promoter was co-infiltrated in all treatments.

### Dual Luciferase Reporter Assay

For protoplast experiments, after overnight incubation, 150 µl were removed from each well and the remaining 50 µl were mixed with 50 µl of 2X Passive Lysis Buffer (Promega), and incubated at 40 rpm for 30 minutes. Fifty microliters of lysate were transferred to white 96-well plates, loaded into a plate reader with automatic injectors (Synergy MX, BioTek), and Firefly (FLUC) and Renilla (REN) luciferase activities were measured using the Dual Luciferase Reporter Assay (Promega) following the addition of 50 µl of each enzyme substrate. Results shown are the average of four technical replicates for the FLUC/REN ratio and each experiment was normalized by its own unrepressed controls.

For agroinfiltration experiments, a 5-mm leaf disc was collected 6 to 8 days after infiltration in a 1.5 mL microcentrifuge tube and ground in liquid nitrogen with the aid of a pestle followed by adding 200 µl of 1X Passive Lysis Buffer. Samples were vortexed briefly and incubated for 30 minutes at 100 rpm. Twenty microliters of lysate were transferred to white 96-well plates and luciferase measurements were carried out as above. For each treatment, three leaves from the same plant were infiltrated in the whole leaf surface and two leaf discs from each leaf were harvested, totaling six replicates per treatment.

### Flavonol Quantification

Flavonol quantification was carried out as described in Chang *et al*. (2020), with modifications. In brief, a 5-mm foliar disc was ground in liquid nitrogen, then 400 µl of 80% methanol (in water) were added, and samples incubated at 4°C for 2 hours. Supernatant was clarified by centrifugation for 5 minutes at 12,000 xg and 300 µl were transferred to a new tube. Then, 100 µl of methanol, 20 µl of AlCl_3_ 10% (w/v), 20 µl of 1 M potassium acetate, and 560 µl of Milli-Q water were added to the sample, followed by incubation at room temperature for 30 minutes. Three hundred microliters were transferred to a 96-well flat bottom clear plate and absorbance at 415 nm was read in a plate reader (Synergy MX, BioTek).

## Results

### Biosensor optimization

We sought to identify bacterial aTFs that could be adapted to sense key metabolites when expressed in plants, and five aTFs were initially selected. Four of them, TtgR, CouR, FapR and PadR, bind to phenylpropanoid-related molecules naringenin, ρ-coumaroyl-CoA, malonyl-CoA and ρ-coumaric acid, respectively, whereas NahR responds to the plant hormone salicylic acid. Naringenin, the first phenylpropanoid pathway intermediate that branches out to flavonoids synthesis, is produced from malonyl-CoA and ρ-coumaroyl-CoA by sequential reactions catalyzed by chalcone synthase (CHS) and chalcone isomerase (CHI). In addition to naringenin, TtgR also binds to structurally similar molecules, such as the flavonol quercetin (Alguel *et al.,* 2007) and the flavone apigenin (Terán *et al.,* 2006). ρ-coumaroyl-CoA is a central metabolite in the phenylpropanoid pathway, which is produced by the enzyme 4-coumarate-CoA ligase (4CL) from ρ-coumarate. Malonyl-CoA is a precursor of both flavonoid and fatty acid biosynthesis and it is produced from acetyl-CoA by acetyl-CoA carboxylase 1 (ACC1), or from malonate by Acyl Activating Enzyme13 (AAE13), also known as malonyl-CoA synthetase (Chen *et al.,* 2011). Lastly, salicylic acid is a plant hormone that triggers immunity responses after pathogen attack (Ding & Ding 2020). All five selected bacterial aTFs function naturally by binding to specific DNA sequences, REs, present in promoters and repressing expression of downstream genes when the aTF’s cognate ligand is absent. If the ligand is present, it associates with the aTF and causes a conformational change in the protein that dissociates it from the REs, activating transcription of downstream genes.

Aiming to adapt these aTFs to function as metabolite sensors in plant cells, we used a similar strategy previously demonstrated (Schaumberg *et al.,* 2016) to engineer the otherwise constitutive CaMV 35S promoter to be controlled by transcription factors. REs bound by each sensor protein were independently inserted at different positions relative to the full-length promoter, namely at the 5’ end, immediately upstream of the TATA-box, and at the 3’ end (Fig. **S1**). To assemble the biosensor devices, two modules are required (Fig. **1a**): the detector module, consisting of a constitutively expressed sensor protein; and the output module, consisting of a reporter gene, firefly luciferase (FLUC), whose expression is controlled by a sensor protein-repressed promoter. To quantitatively test and optimize the function of these biosensor devices, a second reporter, Renilla luciferase (REN), is constitutively expressed to normalize the output module expression. Thus, in the absence of ligand, the aTF represses expression of FLUC, being de-repressed upon ligand binding to the aTF (Fig. **1a**). Repression of these semi-synthetic promoters by their corresponding sensor proteins was tested by transient expression assays in Arabidopsis protoplasts and several rounds of design-build-test (DBT) cycles were required to achieve high levels (>85%) of repression (Figs. **1b-f** and **S2**). The first parameter tested for optimization was the copy number of REs (up to five) inserted in the CaMV 35S promoter (Figs. **S1** and **S2**). FapR and TtgR showed ∼95% repression with one and two RE copies upstream of the TATA-box, respectively (Fig. **1c,f**). PadR, CouR and NahR showed only slight to moderate repression (20 to 70%), regardless of the number of REs (Figs **1b, d, e** and **S2**). Simultaneously with these optimization cycles, we tested whether exogenously applying the ligand to the protoplasts would result in de-repression of the promoters. No de-repression was seen for PadR, CouR and NahR by adding 10 µM of ρ-coumaric acid, ρ-coumaroyl-CoA and salicylic acid, respectively (Fig. **1b, d, e**). For CouR, ρ-coumaroyl-CoA may not cross the plasma membrane (PM) and it may be the reason why no de-repression was detected (Fig. **1b**). FapR is also regulated by a coenzyme A-conjugated metabolite, malonyl-CoA, and therefore not expected to readily cross the PM. Therefore, we first attempted to de-repress the system by expressing two different splice variants of malonyl-CoA synthetase, with or without its substrate, malonate. However, reporter expression was not induced in either case (Fig. **1c**). We also tested expressing ACC1, the other enzyme responsible for producing malonyl-CoA, without success (Fig. **1c**). On the other hand, 10 µM naringenin was capable of de-repressing TtgR (Fig. **1f**). Additional experiments showed that naringenin de-repression is dose dependent, with de-repression fold-change ranging from 2.2- to 6.3-fold when naringenin concentration varies from 1 to 100 µM (Figs. **2** and **S3**). This result opens up the possibility of using TtgR as a real-time sensor for naringenin in plants.

**Fig. 2.**
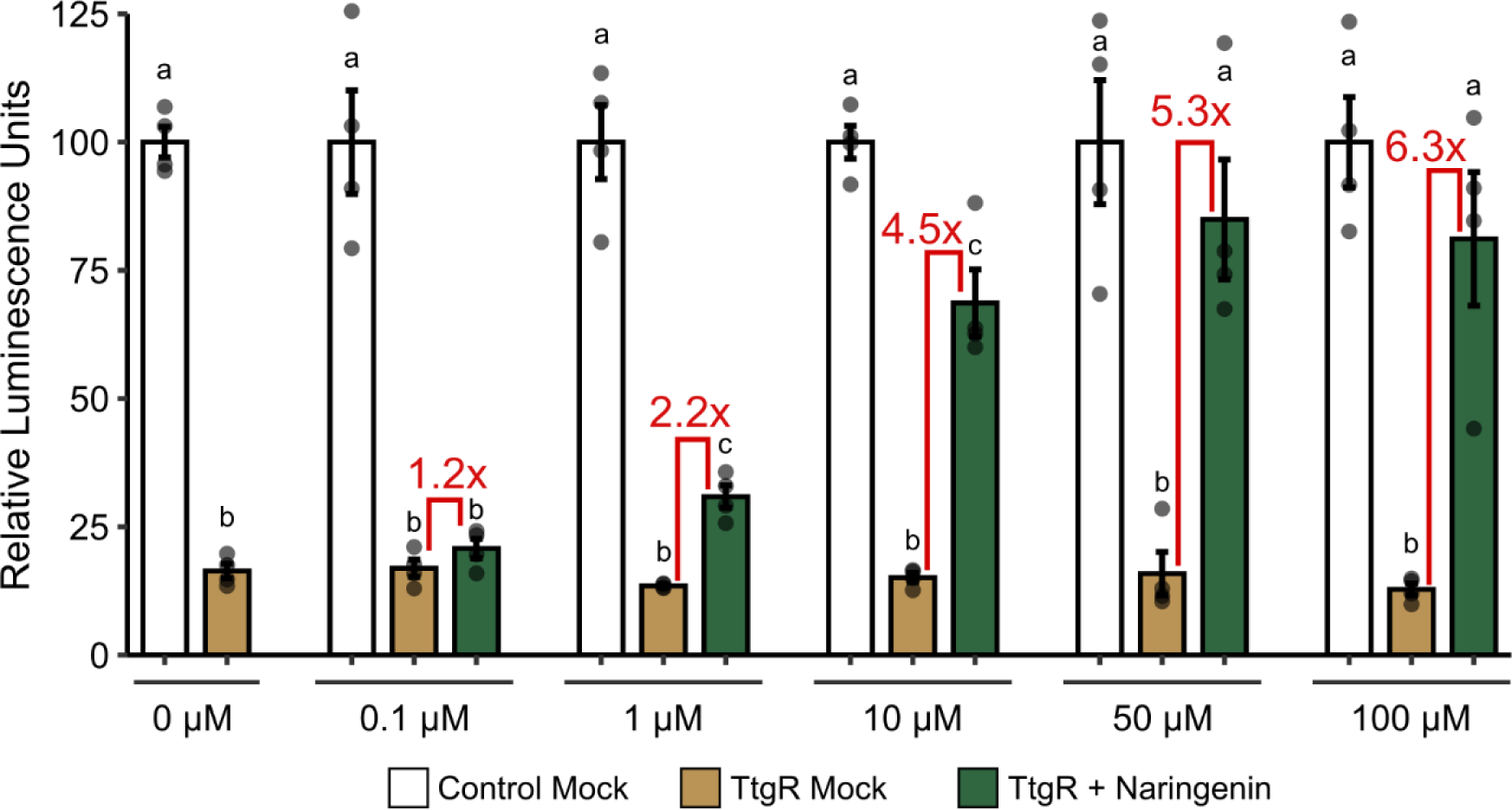
TtgR response curve to naringenin. Transformed protoplasts were treated with different naringenin concentrations or mock treated (DMSO only). Fold induction of expression in de-repressed samples (TtgR + Naringenin) relative to repressed samples (TtgR Mock) are shown. Relative Luminescence Units are the ratio of Firefly (FLUC) and Renilla (REN) Luciferases, normalized by their respective control in each experiment. Error bars = SEM, n = 4, dots represent individual replicates. Statistical significance (*p*<0.05) was determined by One-Way Anova with Tukey’s HSD post hoc test and are depicted by different letters on top of each bar.

Additional DBT cycles for system optimization involved aTF fusions with the Nuclear Localization Signal (NLS), additional repressor domains (SRDX motif), and a protein degradation tag (PEST sequence) (Fig. **S2,** Table **S4**). Only SRDX-CouR showed a significant improvement, achieving 95% repression (Fig. **1b**); however, as no signs of de-repression were seen with the previous experiments, and most likely ρ-coumaroyl-CoA does not cross the protoplast’s PM, we reasoned that further testing SRDX-CouR de-repression in protoplasts with ρ-coumaroyl-CoA would not be relevant. Alternatively, we tested its precursor, ρ-coumaric acid, however, it resulted in no significant de-repression (Fig. **1b**). Summarized results of the best repressible promoter constructs for each aTF are shown in Table **S5.**

### Boolean logic operations using TtgR and FapR in protoplasts

Sensor proteins can be used to provide inputs for genetically encoded circuits built to process information in living cells. Given the remarkably high levels of transcription repression achieved with TtgR and FapR, and the ability to de-repress the TtgR-controlled promoter with naringenin, we sought to design synthetic genetic circuits using these repressors to process information via Boolean logic operations. In digital Boolean logic gates, presence of an input or output is represented by the binary digit 1 (true), whereas absence of these variables is represented by the binary digit 0 (false). The output behavior of a Boolean logic operation, given each combination of the inputs, can thereby be represented by a truth table (Fig. **3**). To build these logic gates, in addition to TtgR, FapR and their respective controlled promoters, we employed a transcription activator consisting of a translational fusion of the yeast Gal4 DBD and the VP64 transactivation domain (Gal4-VP64), which activates a minimal CaMV 35S promoter (−46 – +1) containing five Gal4 REs (*pGal4-min35S*). We also built upon our previously published AND logic gate consisting of a split transcription factor (Anderson *et al.,* 2023), in which VP64 is fused to the P3 hetero-dimerization domain (HD), whereas Gal4 is fused to the P4 cognate domain (HD*). P3-P4 dimerization brings together Gal4 and VP64, fully reestablishing a functional transcriptional factor that activates a *pGal4-min35S* promoter only when both inputs (P3-VP64 and Gal4-P4) are present, characterizing an AND logic operation. The newly built Boolean logic gates were also tested and optimized in transient expression assays in protoplasts, with FLUC used as the quantitative output.

**Fig. 3.**
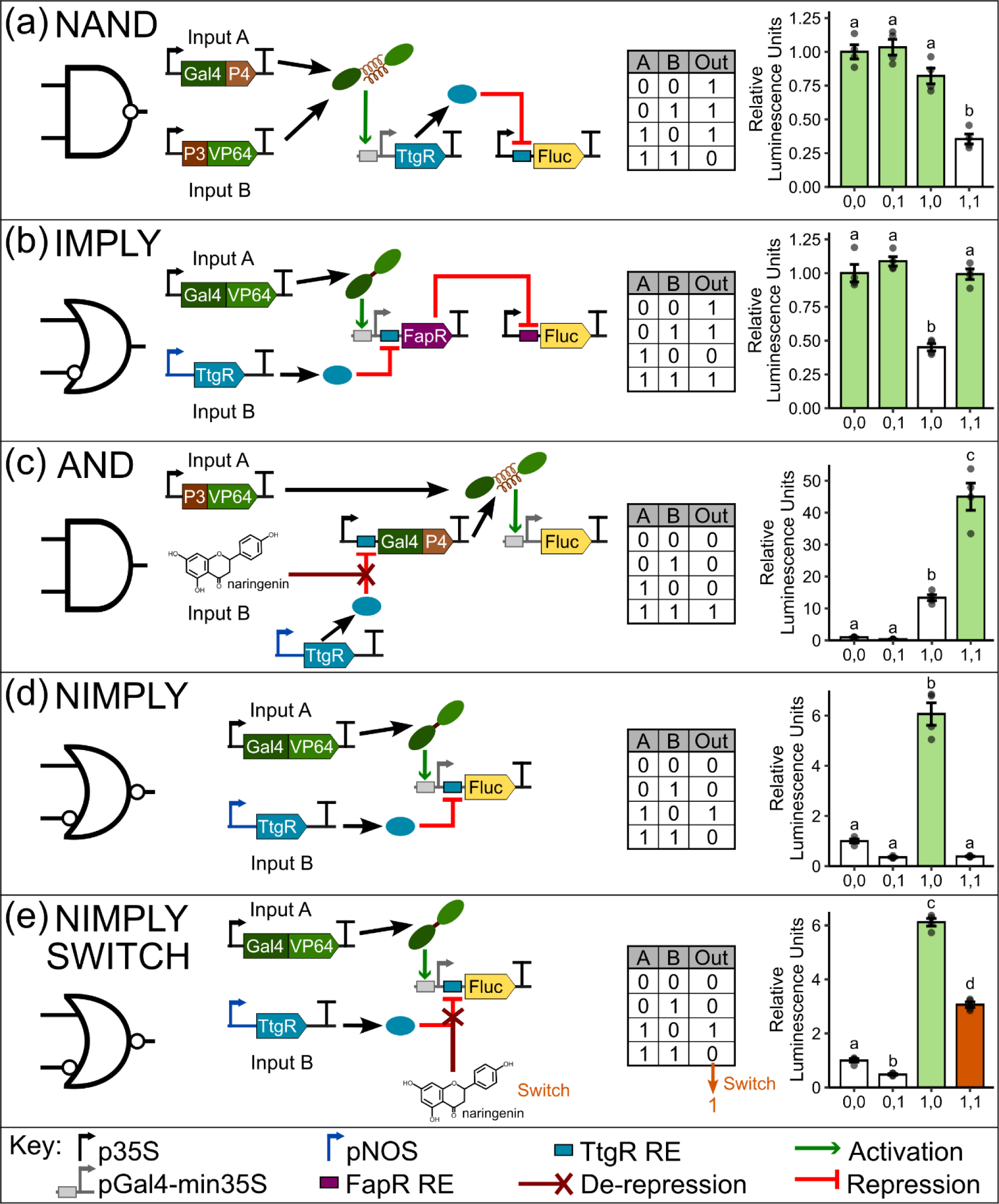
Boolean logic operations in protoplasts. Figure shows the topology of the logic operations NAND (a), IMPLY (b), AND (c), NIMPLY (d) and the NIMPLY switch (e), including inputs and all genetic elements required to build the circuit, as well as the truth table and the diagram for the respective logic. Bar graphs are color-coded to depict expected output expression shown in the truth table as white (output = 0), green (output = 1) and orange (switching output from 0 to 1). Relative Luminescence Units are the ratio of Firefly (FLUC) and Renilla (REN) Luciferases, normalized by their respective control in each experiment. Error bars = SEM, n = 4, dots represent individual replicates. Statistical significance (*p*<0.05) was determined by One-Way Anova with Tukey’s HSD post hoc test and are depicted by different letters on top of each bar. p35S, CaMV 35S promoter; pGal4-min35S, minimal 35S promoter harboring Gal4 binding sequence; pNOS, nopaline synthase promoter; RE, response element.

We first designed a NAND logic gate by adding a NOT (Inverter) operation to our AND gate described above (Anderson *et al.,* 2023). A NAND logic gate has the opposite behavior of an AND gate, with the output being produced with every combination of inputs, except when both inputs are present (Fig. **3a**). To assemble the NAND gate, expression of the naringenin sensor, TtgR, was placed under control of the *pGal4-min35S* promoter, which is activated by reconstitution of the split transcription factor (P3-VP64 + Gal4-P4). Expression of the output was then placed under control of the TtgR-repressible promoter described earlier, essentially inverting the output produced by the AND logic gate (Anderson *et al.,* 2023). Output repression of at least 2.3-fold was seen when both inputs are present, relative to the other input combinations, consistent with a NAND logic operation (Fig. **3a**).

We then tested whether two repressors could be layered to implement an IMPLY logic operation (Fig. **3b**). In a digital IMPLY logic gate, output is produced (true) under every combination of two inputs, except when Input A is present (1) and Input B is absent (0). To implement this logic operation, we placed expression of the reporter gene under control of the FapR repressor. In turn, FapR expression is induced by a direct fusion of the Gal4-VP64 activator and repressed by the TtgR repressor, with the latter two proteins serving as inputs of the IMPLY gate. Thus, output expression is repressed (0) only when FapR is induced by Input A (Gal4-VP64), while at the same time Input B (TtgR) is absent. All other combinations of Inputs A and B should result in FapR being repressed, and thereby the output is produced (Fig. **3b**). We observed at least 2.2-fold repression of the output when Gal4-VP64 was present and TtgR was absent (1,0), relative to the other input combinations, therefore indicating that two layers of repression can be used to design a Boolean logic gate (Fig. **3b**).

We next used naringenin as an input, thereby testing the use of TtgR to integrate metabolite levels into a genetic circuit response. We assembled an AND gate by placing Gal4-P4 expression under control of a TtgR-controlled promoter, so naringenin can be used as one of the inputs, and P3-VP64 as the second input (Fig. **3c**), whereas TtgR is constitutively expressed. Output expression in the resulting AND logic gate is controlled by the *pGal4-min35S* promoter, and thus requires a fully restored transcription factor by dimerization of the P3-P4 pair. However, because Gal4-P4 expression is repressed by TtgR, naringenin is required to de-repress the TtgR-controlled promoter (Fig. **3c**). Although we observed some leaky FLUC expression when only Input A was present (1,0), a 3.3-fold FLUC induction was seen with both inputs present (1,1) (Fig. **3c**). The moderate activation seen with Input A only is likely due to residual expression (∼5%, see Fig. **1f**) from the TtgR-controlled promoter, which allows some expression of Gal4-P4 to pair with P3-VP64 and activate reporter expression (Fig. **3c**).

Finally, we assembled a NIMPLY gate, in which the output response is the reverse of that expected from an IMPLY logic gate. In this NIMPLY gate topology, FLUC expression is controlled by Gal4-VP64 and TtgR (Fig. **3d**), such that only when Gal4-VP64 (Input A) is present, and not TtgR (Input B), output expression is activated. This NIMPLY gate showed at least 6-fold induction with the correct input combination (1,0) (Fig. **3d**). Importantly, in this gate, naringenin can be used as a switch, changing output expression when both inputs are present by de-repressing TtgR function. The naringenin switch successfully achieved a 3-fold induction in the presence of both inputs (Fig. **3e**), without affecting fold change in the other combination of inputs. Again, a lower fold change was expected for the switch compared to the Input A only, because the de-repression with naringenin does not always achieve 100% recovery of the output (Fig. **2**). Nevertheless, this result represents ∼50% of the activation observed with Input A only. Taken together, these results suggest that the metabolite sensors, TtgR and FapR, can be used as efficient repressors to assemble Boolean logic gates in plant protoplasts.

### Boolean logic operations using TtgR and FapR in *N. benthamiana* leaves

After demonstrating successful genetically encoded Boolean logic operations in protoplasts, we used a different transient expression system, Agrobacterium infiltration of *N. benthamiana* leaves, to test whether the same logic operations could be replicated in whole leaves. Unlike the protoplast-based expression system, agroinfiltration allows longer incubation times from infiltration to analysis, and therefore we could test the effect of changing the levels of endogenous metabolites by overexpressing key biosynthetic enzymes and transcription factors. For these experiments, genes encoding specific metabolic enzymes of the pathway or transcription factors that upregulate these enzymes were constitutively expressed from the CaMV 35S promoter and co-infiltrated with plasmids containing sensor proteins and logic gates.

We first tried to de-repress FapR by overexpressing *ACC1*, however, no signs of promoter de-repression could be detected (Fig. **4a**). We also tested if TtgR could be de-repressed in agroinfiltrated leaves by overexpressing enzymes responsible for the synthesis of its ligands, rather than by adding naringenin directly to the cells. In addition to CHS, which catalyzes the conversion of 4-coumaroyl-CoA and malonyl-CoA to naringenin chalcone, leading to increased levels of naringenin, we also tested overexpression of MYB12, a transcription factor that specifically induces flavonol biosynthesis (Mehrtens *et al.,* 2005), as TtgR binds to flavonols such as quercetin (Alguel *et al.,* 2007). Similar to the results obtained in protoplasts, both treatments showed robust de-repression (Fig. **4b**), with CHS leading to 3.4-fold induction of the output (relative to the repressed control with no ligand), whereas MYB12 resulted in full de-repression, with reporter output levels similar to the non-repressed control. As a negative control, we used PadR to show that the de-repression observed with CHS and MYB12 was not an artifact of the agroinfiltration (Fig. **4b**), as no de-repression was seen with PadR. These results indicate that TtgR can respond to changes in the levels of endogenous plant metabolites. To confirm that the de-repression observed in the MYB12 treatment was due to flavonols, we repeated the experiment to measure flavonol accumulation, which showed that overexpression of MYB12 indeed increased flavonol concentration (Fig. **S4**).

**Fig. 4.**
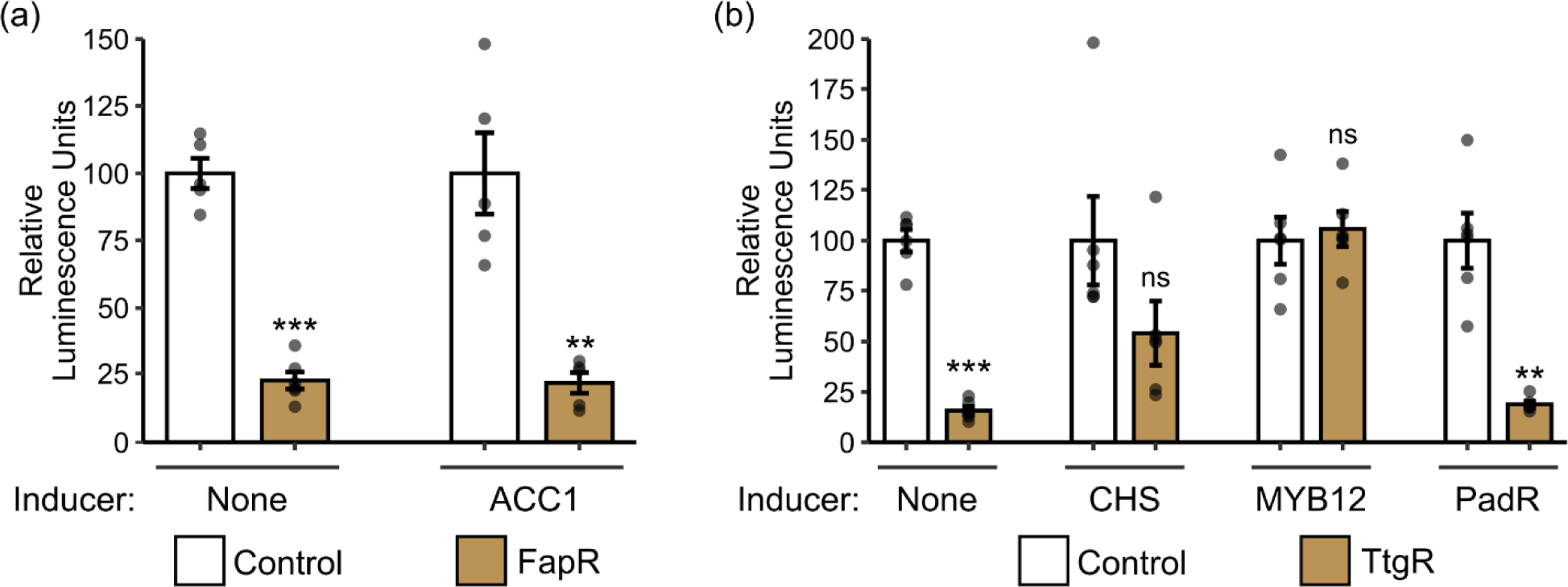
FapR and TtgR transient assay in *N. benthamiana* agroinfiltrated leaves. Transient assay to test the capacity of FapR (a) and TtgR (b) to repress a CaMV 35S promoter harboring their respective RE in agroinfiltrated leaves and to de-repress the system by overexpressing the enzymes or transcription factors responsible for inducing the biosynthesis of their ligands. In (b), PadR is used as negative control for de-repression (inducer). Relative Luminescence Units are the ratio of Firefly (FLUC) and Renilla (REN) Luciferases, normalized by their respective control in each experiment. Error bars = SEM, n = 6, dots represent individual replicates. Statistical significance was determined by 2-tailed Student T-test comparing repressed/induced sample to their respective unrepressed control sample (\**p*<0.05, \*\**p*<0.01 \*\*\**p*<0.001, ns, non-significant). Abbreviations: ACC1, acetyl-CoA carboxylase1; CHS, chalcone synthase.

Next, we sought to further evaluate the use of TtgR in Boolean logic gates that can respond to changes in the levels of endogenous metabolites in *N. benthamiana* leaf tissue. We assembled a similar AND logic gate to the one tested in protoplasts (Fig. **3c**), although here, MYB12 was used as one of the inputs instead of naringenin (Fig. **5a**). Clearly, the increased flavonol levels due to MYB12 overexpression could be detected by TtgR and integrated into an AND logic operation (Fig. **5a**), showing over 15-fold activation of the output only when both inputs are present (1,1). Moreover, a NIMPLY gate was assembled also with the same topology as the one tested in protoplasts (Fig. **3d**), with similar results (Fig. **5b**). When Input A alone is present (1,0), the gate achieved 24-fold induction compared to no inputs (0,0), and 6-fold induction compared to both inputs (1,1) (Fig. **5b**). Also mimicking the NIMPLY gate demonstrated in protoplasts, we introduced a switch, however, in this case we used MYB12 instead of naringenin. As expected, this flavonol switch (Fig. **5c**) successfully increased output expression only when both inputs are present (1,1). Although it did not achieve the same levels of activation compared to Input A only (1,0) (Fig. **5c**), representing ∼50% of its activation, the switch achieved 26-fold induction compared to the No Inputs treatment (0,0). These results demonstrate that the sensor protein TtgR can be used as one of the inputs of a Boolean logic gate, and that changes in leaf flavonol levels can be detected to switch the logic operation.

**Fig. 5.**
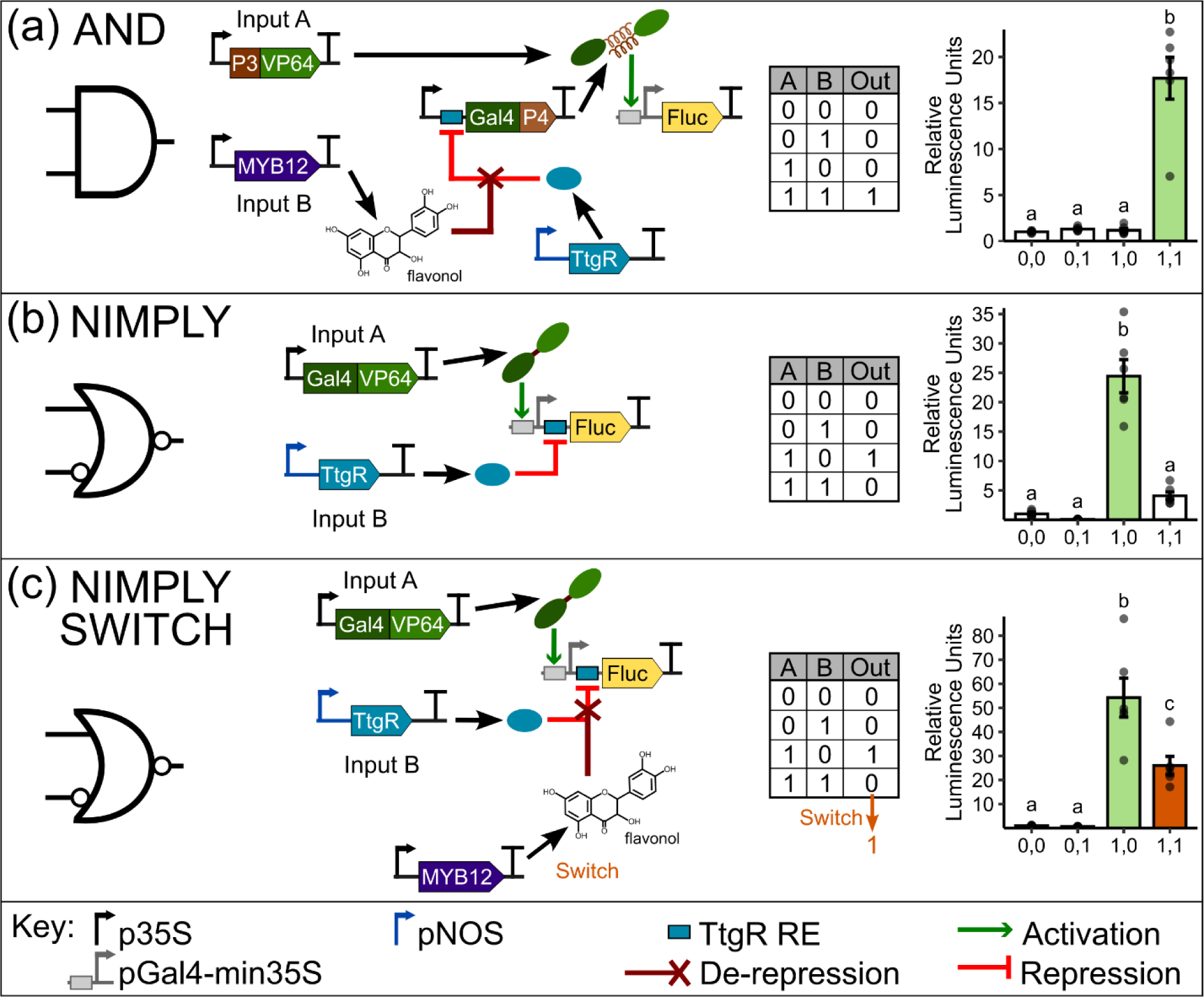
Boolean logic operations in *N. benthamiana* agroinfiltrated leaves. Figure shows the topology of the logic operation AND (a), NIMPLY (b) and the NIMPLY switch (c), including inputs and all genetic elements required to build the genetic circuit, as well as the truth table and the diagram for the respective logic. Bar graphs are color-coded to depict expected output expression as shown in the truth table as white (output = 0), green (output = 1) and orange (switching output from 0 to 1). Relative Luminescence Units are the ratio of Firefly (FLUC) and Renilla (REN) Luciferases, normalized by their respective control in each experiment. Error bars = SEM, n = 6, dots represent individual replicates. Statistical significance (*p*<0.05) was determined by One-Way Anova with Tukey’s HSD post hoc test and are depicted by different letters on top of each bar.

## Discussion

As new challenges emerge with the changing climate, innovative solutions must be introduced to assist us in coping with threats to our food security and public health. Plant synthetic biology can greatly contribute to the next Green Revolution, by engineering crops for a more sustainable agriculture (Kocaoglan *et al.,* 2023; Wang & Demirer 2023; Yang & Reyna-Llorens 2023) and also for bioproduction (Burnett & Burnett 2020; Yang *et al.,* 2022). In the past two decades, efforts in creating cutting-edge approaches in cloning, part standardization, plant transformation, genome engineering, biosensors and genetic circuits (Patron 2020; Vazquez-Vilar *et al.,* 2023; Yang & Reyna-Llorens 2023; Kocaoglan *et al.,* 2023) have profoundly impacted plant synthetic biology. However, many bottlenecks must be addressed before plant synthetic biology-derived products could be made available (Vazquez-Vilar *et al.,* 2023) and, therefore, developing new tools to push the boundaries of plant synthetic biology is essential to unlock its full potential.

In this work, we have shown that bacterial aTFs that bind plant metabolites can be used to control gene expression in plants in a ligand-dependent manner. From five different repressor aTFs tested, two of them could not achieve high levels of repression of our semi-synthetic promoter (Fig. **1**). This can be partially explained by the context of the RE within the CaMV 35S promoter, as the different RE sequences might affect the constitutive promoter strength or aTF binding. A more detailed approach could be implemented by designing fully synthetic promoters (Cai *et al.,* 2020) harboring the REs, and several DBT cycles might be required to test different combinations of cis-regulatory elements to achieve best performing repressor-promoter pairs. Additionally, we showed that only TtgR was able to detect ligand levels, whereas CouR, FapR, PadR, and NahR were not (**Figs 1** and **2**). One possible explanation is that complex molecules, such as CoA conjugates (in the case of FapR and CouR), may not move freely to the nucleus, where the aTF is expected to be localized. However, CouR has been used as ρ-coumaroyl-CoA sensor in yeast (Liu *et al.,* 2022), whereas FapR has been reported as malonyl-CoA sensor in both yeast (Chen *et al.,* 2018) and mammalian cells (Ellis & Wolfgang 2012). Another possibility is that some of these key phenylpropanoid pathway intermediates may not be found freely in the plant cell, as evidence exists that they could be channeled from one enzymatic step to the next in the pathway through enzyme complexes called metabolons (Biala & Jasinski 2018). Still, malonyl-CoA is a substrate for different pathways, such as the lignin and flavonoid branches of the phenylpropanoid pathway, as well as fatty acid elongation. Hence, this molecule is likely to be available in the cytoplasm to be used as a substrate by different enzymes, although cooperation of different metabolons cannot be excluded (Biala & Jasinski 2018). Alternatively, physiological ligand concentrations might fall below the dynamic range of the sensor, or the applied concentrations might be too high, making the system unable to respond. To overcome these constraints, strategies such as the introduction of mutations (Meyer *et al.,* 2019; Beltrán *et al.,* 2022) to evolve the biosensors and improve their dynamic range or to change ligand specificity could be employed to broaden their use.

Genetically-encoded biosensors have been used for several applications in plants. Much attention has been given to biosensors capable of detecting and reporting *in planta* the spatio-temporal levels of specific molecules, such as sugars and hormones (reviewed by Walia *et al.,* 2018; Wright & Nemhauser 2019; Vazquez-Vilar *et al.,* 2023). Other studies have used biosensors to engineer expression systems inducible by exogenous substances or to detect molecules in the environment (Dugdale *et al.,* 2013; Garcia-Perez *et al.,* 2022), working as basic on-off transcriptional switches. Nonetheless, only a limited number of studies have employed aTFs, or other synthetic biology devices, to integrate ligand levels into a more complex response, which is key for innovative plant improvements (Vazquez-Vilar *et al.,* 2023). Antunes *et al*. (2011) engineered a system with a bacterial periplasmic binding protein that detects the explosive TNT, and rewired its downstream signaling to activate a kinase that ultimately activates chlorophyll degradation, causing plant de-greening in the presence of TNT. Iacopino *et al*. (2019) built an oxygen sensing device in plants based on animal hypoxia signaling components, functioning as a tool to tune gene expression based on oxygen availability, which could help the plant to cope with flooding, for example. Finally, Khakhar et al. (2018) developed synthetic hormone responsive transcription factors that could re-program plant development by changing the strength of auxin response mechanisms to feedback on the expression of its transporter PIN-FORMED1, hence altering plant shoot architecture. In the present work, we have shown for the first time an aTF capable of sensing endogenous metabolites from specialized metabolism in plants and integrating their levels into a genetic circuit that computes information using Boolean logic operations. These results open new opportunities for plant metabolism customization, with the possibility of designing regulatory networks related to bioproduction or enhanced plant defense.

Complex circuits controlled by biosensors have been demonstrated to improve value-added microbial chemical production of naringenin (Zhou *et al.,* 2021) and fatty acids (Xu *et al.,* 2014). In these circuits, the levels of metabolic intermediates could be measured in real time, triggering changes in circuit behavior to achieve dynamic control of the pathway. The work described here shows that this technology can also be applied in plants, and it can be further combined with other approaches for precise control of metabolic pathways in plants as well. For example, TtgR could be incorporated into a genetic circuit as an additional layer of control to the work of Selma *et al*. (2022), where selective activation of genes in the phenylpropanoid pathway was carried out to customize flavonoid profile. TtgR could be used to fine tune expression and reduce negative feedback mechanisms, increasing yields while reducing fitness penalty. This strategy enables the organism to adjust bioproduction “on-the-fly” according to environmental cues and intracellular changes, without external interference (Gao *et al.,* 2019). Nonetheless, some optimization may be required, including expanding the biosensors repertoire with other characterized aTFs that sense different plant metabolites (Garagounis *et al.,* 2021; Ferreira & Antunes 2021), or by building chimeric aTFs through DBD and LBD swapping (Dimas *et al.,* 2019) to create new ligand-RE pairs. The additional aTFs could thus sense several key intermediates of the pathway, allowing logic gates to be layered to compute multiple (>2) inputs, and leading to the development of more robust circuits, *i.e.*, with a lower probability of unwanted activation of the output. Even though we could not test multiple metabolite as inputs, our IMPLY gate (Fig. **3b**) shows that at least two aTFs can be successfully layered to build a logic operation.

The plant phenylpropanoid pathway produces a large number of compounds with important properties, from antimicrobial to fragrances (Neelam *et al.,* 2020). A rich diversity of phenylpropanoids is synthesized by different plant species, with a variety of specialized enzymes that could be tapped for engineering more tractable plant species for high production yields of these useful compounds. Regulation of this pathway is complex and maintained at many levels, including transcriptional, translational and post-translational, and even by spatial availability of intermediates in different cell compartments. Despite this regulatory complexity, intentional redirection of metabolite flux to different branches of this pathway has been previously demonstrated with simple genetic alterations (*e.g.*, mutations, over-expression) (Mayer *et al.,* 2001; Mir Derikvand *et al.,* 2008; Lanot *et al.,* 2008; Menard *et al.,* 2022). However, these permanent modifications tend to result in reduced plant fitness, with detrimental effects on plant growth and resistance to pests and pathogens. Genetic circuits that allow dynamic control of a metabolic pathway in response to metabolite availability could overcome these limitations and result in plants that use resources more efficiently, and as a result are ideal chassis for bioproduction. Such genetically encoded synthetic circuits capable of integrating multiple signal using logic operations have only recently started to become a reality in plants, bestowing scientists the ability to re-program and re-shape plant morphology and physiology. For example, synthetic circuits have been designed with Boolean logic gates to re-program root development (Brophy *et al.,* 2022), and to integrate stress signals into a coordinated response (Anderson *et al.,* 2023). In this study, we showed that AND, NAND, IMPLY and NIMPLY Boolean logic operations can be engineered into plants using aTFs as inputs for the circuit (**Figs 3** and **5**). Moreover, these metabolites can serve as switches to dynamically change the gate logic (**Figs 3e** and **5c**). The flexibility afforded by these logic gates enhances our ability to fine-tune and optimize plant responses, ultimately improving the overall efficacy of synthetic biology applications.

## Acknowledgements

This work was supported by startup funds from the Department of Biological Sciences and the BioDiscovery Institute at the University of North Texas to M.S.A. We thank Camila Lima and Ruhbani Jarral for help in processing samples.

## Author Contributions

M.S.A. and S.S.F. conceived the study; S.S.F performed the experiments; M.S.A. and S.S.F designed the experiments, analyzed the results and prepared the manuscript.

## Data availability

The data that supports the findings of this study are available within the manuscript and its supplementary information.

