## Supporting Information for "Genetically encoded Boolean logic operators to sense and integrate phenylpropanoid metabolite levels in plants"

The following Supporting Information is available for this article:

**Fig. S1** Schematic representation of semi-synthetic promoters harboring the response element for each aTF.

**Fig. S2** Additional optimization tests for different promoter constructs and aTF fusions.

**Fig. S3** Independent experiment showing TtgR response curve to naringenin in protoplasts.

**Fig. S4** Flavonol quantification in agroinfiltrated leaves.

**Table S1** Nucleotide sequences of the aTFs used in this study

**Table S2** Nucleotide sequences of the semi-synthetic CaMV 35S promoters harboring the response element (shown in bold) of each aTF used in this study.

**Table S3** Primers used in this study.

**Table S4** Nucleotide sequences of additional domains fused to aTFs, promoter and other protein fusions not built in this work.

**Table S5** Promoter constructs that showed the highest repression level for each respective aTF.

**Fig. S1** Schematic representation of semi-synthetic promoters harboring the response element for each aTF. Response elements were located at the 5’, upstream TATA box or at the 3’ of the CaMV 35S promoter. Multiple combinations of number of response elements were tested as shown below. Note that not all combinations were tested for all aTFs. Sequences for all promoter constructs are shown in Table **S2**. FLUC, Firefly Luciferase coding sequence.

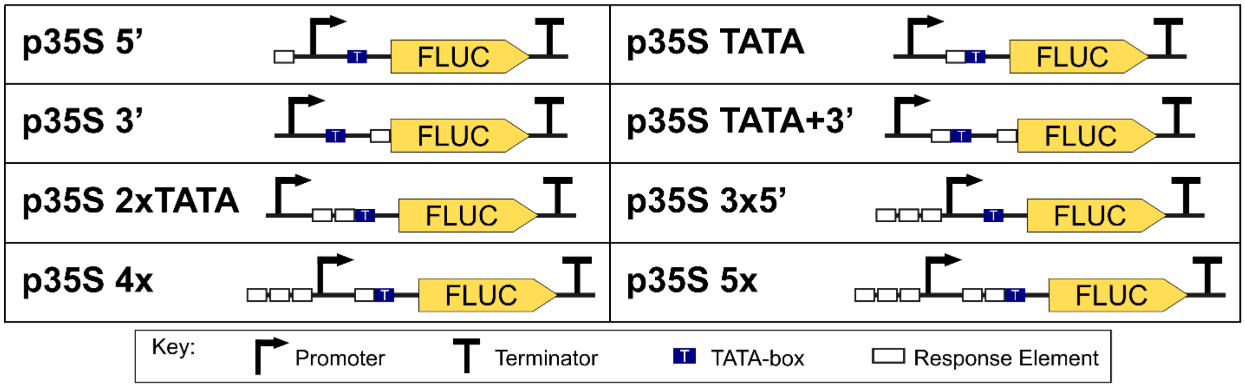

**Fig. S2** Additional optimization tests for different promoter constructs and aTF fusions. For each aTF responsive promoter, the construct with highest repression level was further tested with different aTF fusions. Relative Luminescence Units are the ratio of Firefly (FLUC) and Renilla (REN) Luciferases, normalized by their respective control in each experiment. Error bars = SEM, n = 4, dots represent individual replicates. Statistical significance was determined by 2-tailed Student T-test comparing repressed/induced sample to their respective unrepressed control sample (*p<0.05, **p<0.01 ***p<0.001, ns, non-significant). SRDX, EAR-repression domain; PEST, degradation tag; NLS, nuclear localization signal.

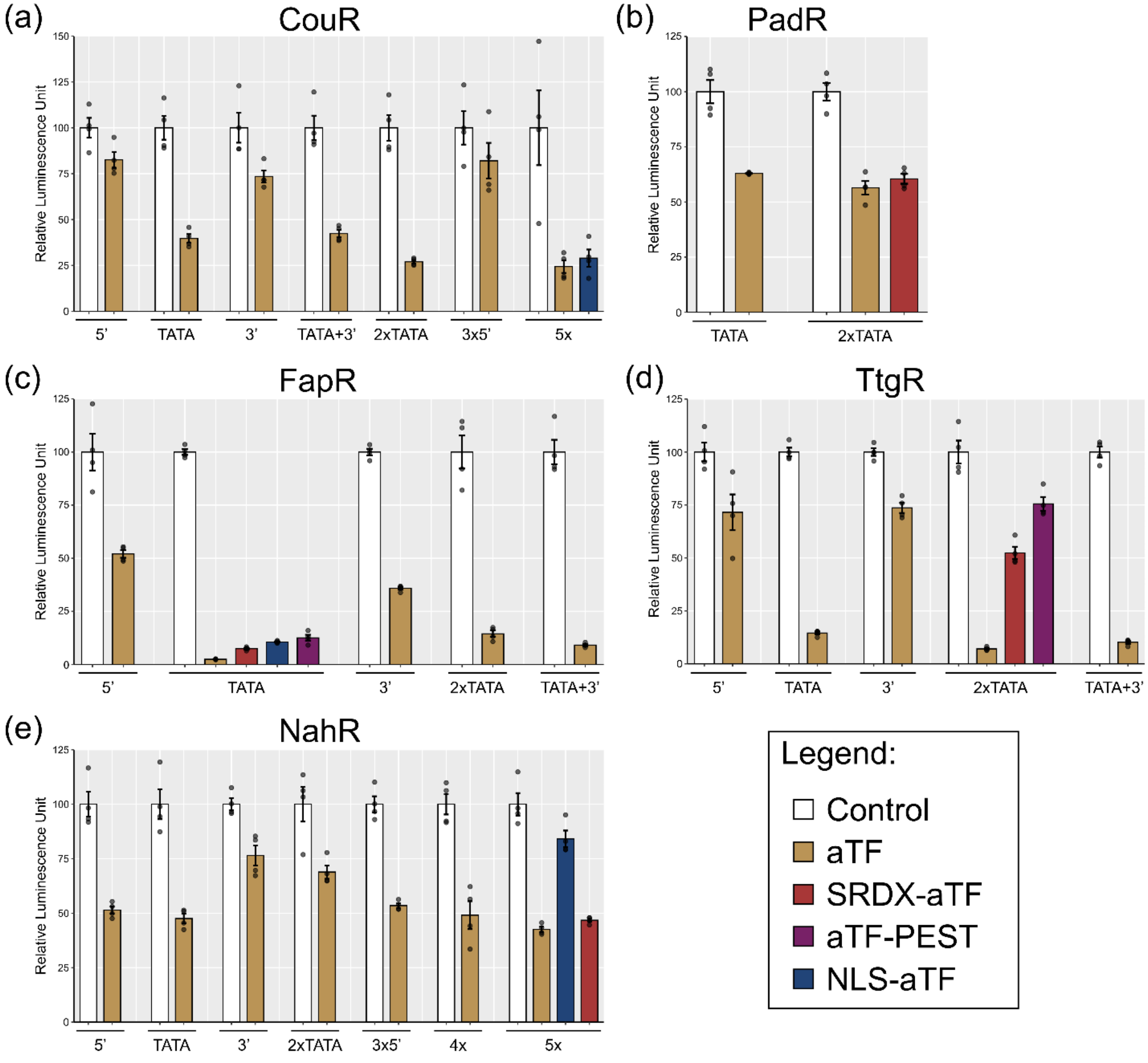

**Fig. S3** Independent experiment showing TtgR response curve to naringenin in protoplasts. De-repression induced by increasing concentrations of naringenin is shown on the left y-axis, whereas fold change induction of de-repressed samples (TtgR + Naringenin) relative to repressed samples (TtgR 0 µM, treated with DMSO only) are shown in the right y-axis (red line). Relative Luminescence Units are the ratio of Firefly (FLUC) and Renilla (REN) Luciferases, normalized by their respective control in each experiment. Error bars = SEM, n = 4, dots represent individual replicates. Statistical significance was determined by 2-tailed Student T-test comparing repressed/induced sample to their respective unrepressed control sample (*p<0.05, **p<0.01 ***p<0.001, ns, non-significant).

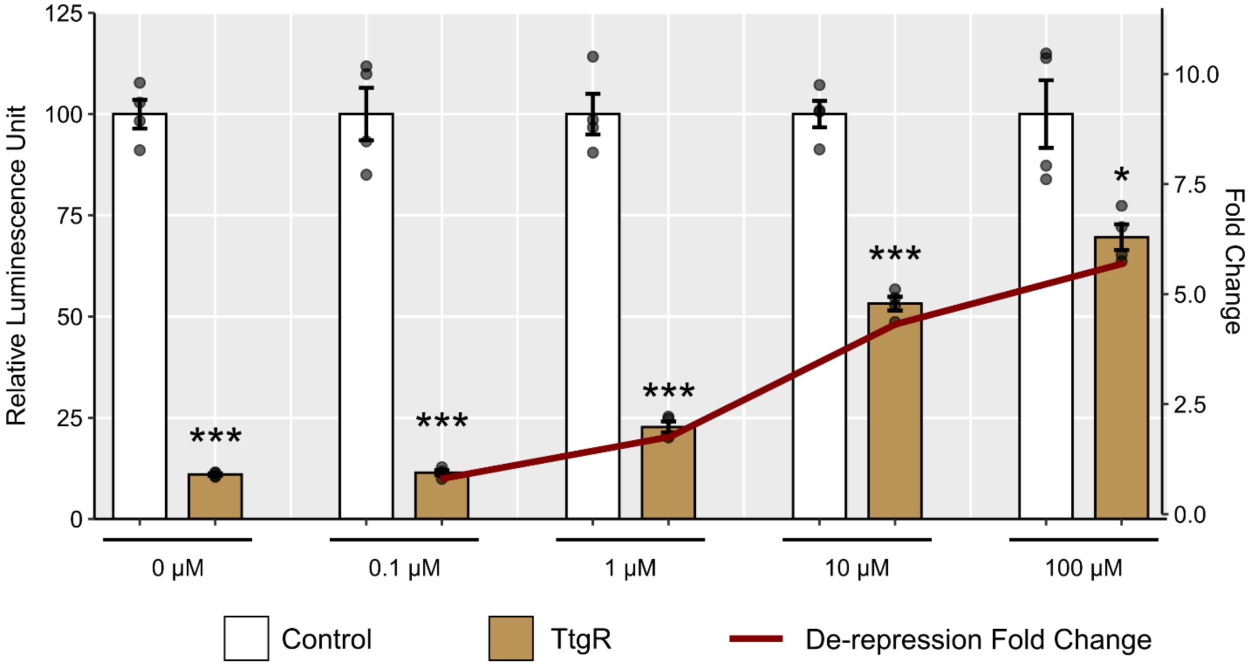

**Fig. S4** Flavonol quantification in agroinfiltrated leaves. Flavonol quantities are determined by absorbance of the phenolic extract at 415 nm. Error bars = SEM, n = 6, dots represent individual replicates. Statistical significance ($p<0.05$) was determined by One-Way Anova with Tukey's HSD post hoc test and are depicted by different letters on top of each bar. CHS, chalcone synthase.

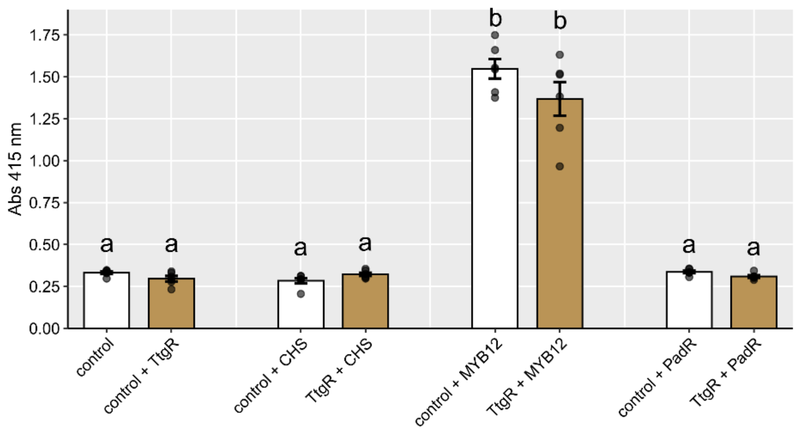


**Table S1** Nucleotide sequences of the aTFs used in this study.

| | |
|---|---|
| CouR | Otani *et al.*, 2016 |
| ATGTCCGGGAGTATGACTAGCTCTAATCGAATCACATCACCCGCAATGACAGCGA<br>GTAAAACCGCCGCCGTAGCAAAGCCGACCCGTGCCGGACGTAAAGCTCCAGCGGT<br>AGAAACAGCACCGGAAGCATCCGAGCTCAAAATGGGTGAGTTATCAGAATTGCTC<br>GGTTATGCGCTTAAGAGGGCACAACTACGTGTCTTCGAGGATTTTTACATTGTGT<br>AGCTCCTGTGCAGCTTACTCCAGCCCAATTCAGCGTACTCCTCCTACTGGACGCTA<br>ATCCCGGAAGGAACCAGACGGAAATAGCGACCACGCTTGGAATATTGAGGCCAA<br>ATTTTGTAGCTATGCTAGATGCCCTAGAGGGAAGGGGTTTATGCGTCAGAACCCG<br>ATCCCCAAGTGACCGAAGGTCCCATATTCTAATGCTTACTGATAAGGGCAGGGCA<br>ACTCTAGCAAGGGCAAAAAAGCTGGTGGCCACTAGGCACGAAGATAGGCTCACG<br>GAGCTCCTGGGGAGGGACAACCGGGACGCCCTGCTTTCCATGTTGGCAACTATAG<br>CAAGGGAATTCTAA | |
| FapR | Xu *et al.*, 2014 |
| ATGCGTCGAAACAAAAGGGAAAGGCAAGAATTGCTCCAACAAACAATACAGGCA<br>ACGCCATTTATCACGGACGAGGAGCTAGCAGGGAAGTTTGGTGTCTCCATTCAAA<br>CCATTCGACTAGATCGTTTAGAACTCTCTATACCAGAATTGCGAGAACGAATCAA<br>AAACGTAGCTGAAAAGACCCTGGAGGACGAGGTAAAATCCCTCTCTTTGGATGAA<br>GTGATCGGCGAGATAATCGACCTGGAACTGGATGATCAAGCAATCTCTATCCTCG<br>AAATTAAACAAGAACACGTCTTTAGTAGGAATCAGATTGCGCGTGGGCACCACCT<br>ATTCGCTCAGGCAAATTCTCTTGCCGTAGCGGTGATCGACGACGAGCTGGCTTTAA<br>CTGCATCTGCTGATATAAGATTCACCCGTCAAGTGAAGCAGGGTGAACGTGTGGT | |

<table>
<tr><td colspan="2">CGCGAAGGCTAAAGTTACCGCCGTTGAAAAAGAAAAAGGCCGTACTGTCGTCGAG<br>GTCAACTCTTATGTGGGGGAGGAAATCGTGTTCTCTGGAAGATTTGATATGTATAG<br>AAGTAAACATTCCTAA</td></tr>
<tr><td>NahR</td><td>Meyer et al., 2019</td></tr>
<tr><td colspan="2">ATGGAATTACGAGACTTAGACTTGAACCTTTTGGTCGTTTTCAACCAGCTCTTAGT<br>GGATAGGAGGGTTAGTGTTACTGCGGAAAACTTGGGACTCACCCAACCAGCAGTT<br>AGTAACGCGTTGAAAAGGCTGAGAACTAGTTTGCAAGACCCTCTCTTTGTCCGTAC<br>ACACCAAGGGATGGAGCCAACCCCGTACGCGGCTCACCTCGCAGAGCACGTAAC<br>AAGCGCCATGCATGCACTTCGTAACGCGTTACAACACCATGAATCCTTCGATCCCC<br>TAACTTCAGAGCGAACCTTCACTCTGGCCATGACTGATATTGGGGAGATATACTTT<br>ATGCCGCGTCTTATGGATGCTTTAGCTCATCAAGCACCAAACTGCGTAATCTCAAC<br>GGTCAGGGATTCATCTATGAGCCTGATGCAGGCTCTACAAAACGGTACGGTGGAT<br>CTTGCAGTCGGCCTTCTACCGAACCTCCAAACTGGCTTTTTCCAACGAAGGCTTCT<br>TCAAAACCATTACGTGTGCTTATGCCGTAAAGACCACCCTGTGACGCGTGAGCCA<br>CTAACACTAGAGAGATTCTGCAGCTATGGGCATGTGAGGGTGATCGCGGCGGGGA<br>CGGGTCATGGAGAAGTAGATACTTACATGACGAGGGTCGGTATCAGGCGTGACAT<br>TCGACTCGAAGTGCCGCATTTTGCCGCTGTGGGTCACATCCTCCAACGTACAGATT<br>TACTGGCGACCGTTCCCATTTGCCTGGCTGACTGTTGTGTCGAGCCATTTGGATTA<br>TCTGCCCTACCACATCCCGTCGTGCTACCGGAAATTGCCATAAATATGTTCTGGCA<br>TGCAAAATATCACAAGGACCTAGCTAATATCTGGCTGAGACAACTTATGTTCGAC<br>TTATTCACGGATTAA</td></tr>
<tr><td>PadR</td><td>Park et al., 2017</td></tr>
<tr><td colspan="2">ATGAGAGTGTTAAAATATGCAATCCTACGTTTGCTGAGGAAAGGGGAACTAAGTG<br>GGTACGATATAACATCATACTTCAAAGAAGAATTAGGTCAGTTCTGGAGCGCCAA<br>GCACTCCCAGATATATCCTGAGCTAAAGAAACTGACAGATGAAGGTTTTATCACG<br>TTCCGAACTACGATTCAGGGAACCAAGTTGGAGAAGAAGATGTATACTCTTACGG<br>ATTCCGGAAAACAAGAGTTGCACGACTGGTTAATCCGACATCAACCCATCCCGGA<br>AACGGTCAAAGACGAGTTCATGCTTAAGGCTTATTTTATCTCTTGCCTTAGCCGTC<br>AGGAAGCCAGTGATTTGTTTAAAGACCAACTCCAAAAAAGGCAAGCGAAGCTTTC<br>AGATTTACAGGGGAGTTATGAGAAGTTAATGGCTTCTGCTGAACCTATGTCTTTCT<br>CAAGTCCGGATTTCGGACATTACCTTGTCCTGACGAAAGCTTTAGAGCGTGAAAA<br>AAACTACGTGTCATGGCTGGAGTCCATTTTGGCGATGATAGATAAGGACTGA</td></tr>
<tr><td>TtgR</td><td>Meyer et al., 2019</td></tr>
<tr><td colspan="2">ATGGTAAGACGTACCAAAGAGGAAGCTCAGGAGACTCGAGCACAGATAATTGAG<br>GCTGCGGAGCGTGCGTTTTATAAAAGGGGCGTAGCCCGAACAACCCTCGCGGACA<br>TAGCGGAGTTAGCCGGAGTAACTCGAGGAGCGATATATTGGCACTTCAACAATAA<br>GGCTGAACTAGTGCAGGCACTTCTCGACAGTTTGCATGAAACCCATGATCACCTG<br>GCAAGGGCAAGCGAGAGTGAAGACGAGGTGGACCCGCTTGGCTGCATGCGTAAG<br>CTTCTCCTACAAGTGTTCAATGAATTAGTTCTCGATGCAAGAACACGAAGAATAA<br>ACGAAATTCTCCATCATAAATGCGAGTTCACAGACGACATGTGCGAGATAAGACA<br>ACAACACCAGTCCGCAGTATTGGACTGTCATAAAGGAATCACCCTAACCCTCGCT<br>AACGTGGTACGTCGAGGACAACTGCCTGGTGAATTGGACGCTGAACGTGCGGCGG<br>TTGCCATGTTCGCGTATGTGGACGGACTAATTCGACGATGGCTCCTCCTTCCCGAT<br>TCCGTTGACCTCCTCGGCGACGTGGAAAAGTGGGTAGACACCGGCTTAGACATGT<br>TGAGGTTGTCTCCTGCTTTACGAAAGTAA</td></tr>
</table>

**Table S2** Nucleotide sequences of the semi-synthetic CaMV 35S promoters harboring the response element (shown in bold) of each aTF used in this study. TATA-box is highlighted as underlined. Response element sequences were retrieved from the reference as indicated below.

| CouR | Otani *et al.,* 2016; Cogan *et al.,* 2018 |
|---|---|
| p35S 5’ | **TTGTTATACTCTATAACTATTCTGCACAG**TGAGACTTTTCAACAAA<br>GGGTAATATCGGGAAACCTCCTCGGATTCCATTGCCCAGCTATCTGT<br>CACTTCATCAAAAGGACAGTAGAAAAGGAAGGTGGCACCTACAAAT<br>GCCATCATTGCGATAAAGGAAAGGCTATCGTTCAAGATGCCCCTGC<br>CGACAGTGGTCCCAAAGATGGACCCCCACCCACGAGGAGCATCGTG<br>GAAAAAGAAGACGTTCCAACCACGTCTTCAAAGCAAGTGGATTGAT<br>GTGATATCTCCACTGACGTAAGGGATGACGCACAATCCCACTATCCT<br>TCGCAAGACCCTTCCTCTATATAAGGAAGTTCATTTCATTTGGAGAG<br>GACTCCGGTATTTTTACAACAATTACCACAACAAAACAAACAACAA<br>ACAACATTACAATTTACTATTCTAGTCGAAA |
| p35S TATA | TGAGACTTTTCAACAAAGGGTAATATCGGGAAACCTCCTCGGATTCC<br>ATTGCCCAGCTATCTGTCACTTCATCAAAAGGACAGTAGAAAAGGA<br>AGGTGGCACCTACAAATGCCATCATTGCGATAAAGGAAAGGCTATC<br>GTTCAAGATGCCCCTGCCGACAGTGGTCCCAAAGATGGACCCCCAC<br>CCACGAGGAGCATCGTGGAAAAAGAAGACGTTCCAACCACGTCTTC<br>AAAGCAAGTGGATTGATGTGATATCTCCACTGACGTAAGGGATGAC<br>GCACAATCCCACTATCCTTCGCAAGACCCTTCCTC**TTGTTATACTCT**<br>**ATAACTATTCTGCACAG**TATATAAGGAAGTTCATTTCATTTGGAGA<br>GGACTCCGGTATTTTTACAACAATTACCACAACAAAACAAACAACA<br>AACAACATTACAATTTACTATTCTAGTCGAAA |
| p35S 3’ | TGAGACTTTTCAACAAAGGGTAATATCGGGAAACCTCCTCGGATTCC<br>ATTGCCCAGCTATCTGTCACTTCATCAAAAGGACAGTAGAAAAGGA<br>AGGTGGCACCTACAAATGCCATCATTGCGATAAAGGAAAGGCTATC<br>GTTCAAGATGCCCCTGCCGACAGTGGTCCCAAAGATGGACCCCCAC<br>CCACGAGGAGCATCGTGGAAAAAGAAGACGTTCCAACCACGTCTTC<br>AAAGCAAGTGGATTGATGTGATATCTCCACTGACGTAAGGGATGAC<br>GCACAATCCCACTATCCTTCGCAAGACCCTTCCTCTATATAAGGAAG<br>TTCATTTCATTTGGAGAGGACTCCGGTATTTTTACAACAATTACCAC<br>AACAAAACAAACAACAAACAACATTACAATTTACTATTCTAGTCGA<br>A**TTGTTATACTCTATAACTATTCTGCACAG** |
| p35S TATA+3’ | TGAGACTTTTCAACAAAGGGTAATATCGGGAAACCTCCTCGGATTCC<br>ATTGCCCAGCTATCTGTCACTTCATCAAAAGGACAGTAGAAAAGGA<br>AGGTGGCACCTACAAATGCCATCATTGCGATAAAGGAAAGGCTATC<br>GTTCAAGATGCCCCTGCCGACAGTGGTCCCAAAGATGGACCCCCAC<br>CCACGAGGAGCATCGTGGAAAAAGAAGACGTTCCAACCACGTCTTC<br>AAAGCAAGTGGATTGATGTGATATCTCCACTGACGTAAGGGATGAC<br>GCACAATCCCACTATCCTTCGCAAGACCCTTCCTC**TTGTTATACTCT**<br>**ATAACTATTCTGCACAG**TATATAAGGAAGTTCATTTCATTTGGAGA<br>GGACTCCGGTATTTTTACAACAATTACCACAACAAAACAAACAACA<br>AACAACATTACAATTTACTATTCTAGTCGAA**TTGTTATACTCTATA**<br>**ACTATTCTGCACAG**A |

| | |
|---|---|
| p35S 2xTATA | TGAGACTTTTCAACAAAGGGTAATATCGGGAAACCTCCTCGGATTCC<br>ATTGCCCAGCTATCTGTCACTTCATCAAAAGGACAGTAGAAAAGGA<br>AGGTGGCACCTACAAATGCCATCATTGCGATAAAGGAAAGGCTATC<br>GTTCAAGATGCCCCTGCCGACAGTGGTCCCAAAGATGGACCCCCAC<br>CCACGAGGAGCATCGTGGAAAAAGAAGACGTTCCAACCACGTCTTC<br>AAAGCAAGTGGATTGATGTGATATCTCCACTGACGTAAGGGATGAC<br>GCACAATCCCACTATCCTTCGCAAGACCCTTCCTC**TTGTTATACTCT**<br>**ATAACTATACTGTGCTG**GAGGCACGAC**TTGTTATACTCTATAACT**<br>**ATTCTGCACAG**TATATAAGGAAGTTCATTTCATTTGGAGAGGACTC<br>CGGTATTTTTACAACAATTACCACAACAAAACAAACAACAAACAAC<br>ATTACAATTTACTATTCTAGTCGAAA |
| p35S 3x5’ | **TTGTTATACTCTATAACTATACTGTGCTG**TAGGCACGTA**TTGTTA**<br>**TACTCTATAACTATATTGCACAG**GCTCAGAACG**TTGTTATACTCT**<br>**ATAACTATTCTGCACAG**TGAGACTTTTCAACAAAGGGTAATATCGG<br>GAAACCTCCTCGGATTCCATTGCCCAGCTATCTGTCACTTCATCAAA<br>AGGACAGTAGAAAAGGAAGGTGGCACCTACAAATGCCATCATTGCG<br>ATAAAGGAAAGGCTATCGTTCAAGATGCCCCTGCCGACAGTGGTCC<br>CAAAGATGGACCCCCACCCACGAGGAGCATCGTGGAAAAAGAAGA<br>CGTTCCAACCACGTCTTCAAAGCAAGTGGATTGATGTGATATCTCCA<br>CTGACGTAAGGGATGACGCACAATCCCACTATCCTTCGCAAGACCCT<br>TCCTCTATATAAGGAAGTTCATTTCATTTGGAGAGGACTCCGGTATT<br>TTTACAACAATTACCACAACAAAACAAACAACAAACAACATTACAA<br>TTTACTATTCTAGTCGAAA |
| p35S 5x | **TTGTTATACTCTATAACTATACTGTGCTG**TAGGCACGTA**TTGTTA**<br>**TACTCTATAACTATATTGCACAG**GCTCAGAACG**TTGTTATACTCT**<br>**ATAACTATTCTGCACAG**TGAGACTTTTCAACAAAGGGTAATATCGG<br>GAAACCTCCTCGGATTCCATTGCCCAGCTATCTGTCACTTCATCAAA<br>AGGACAGTAGAAAAGGAAGGTGGCACCTACAAATGCCATCATTGCG<br>ATAAAGGAAAGGCTATCGTTCAAGATGCCCCTGCCGACAGTGGTCC<br>CAAAGATGGACCCCCACCCACGAGGAGCATCGTGGAAAAAGAAGA<br>CGTTCCAACCACGTCTTCAAAGCAAGTGGATTGATGTGATATCTCCA<br>CTGACGTAAGGGATGACGCACAATCCCACTATCCTTCGCAAGACCCT<br>TCCTC**TTGTTATACTCTATAACTATACTGTGCTG**GAGGCACGAC**TT**<br>**GTTATACTCTATAACTATTCTGCACAG**TATATAAGGAAGTTCATTT<br>CATTTGGAGAGGACTCCGGTATTTTTACAACAATTACCACAACAAAA<br>CAAACAACAAACAACATTACAATTTACTATTCTAGTCGAAA |
| FapR | Schujman *et al.,* 2003; David *et al.,* 2016 |
| p35S 5’ | **AATTATATACTACTATTAGTACCTAGTCTTAATT**TGAGACTTTTCA<br>ACAAAGGGTAATATCGGGAAACCTCCTCGGATTCCATTGCCCAGCT<br>ATCTGTCACTTCATCAAAAGGACAGTAGAAAAGGAAGGTGGCACCT<br>ACAAATGCCATCATTGCGATAAAGGAAAGGCTATCGTTCAAGATGC<br>CCCTGCCGACAGTGGTCCCAAAGATGGACCCCCACCCACGAGGAGC<br>ATCGTGGAAAAAGAAGACGTTCCAACCACGTCTTCAAAGCAAGTGG<br>ATTGATGTGATATCTCCACTGACGTAAGGGATGACGCACAATCCCAC<br>TATCCTTCGCAAGACCCTTCCTCTATATAAGGAAGTTCATTTCATTTG |

| | |
|---|---|
| | GAGAGGACTCCGGTATTTTTACAACAATTACCACAACAAAACAAACAACAAACAACATTACAATTTACTATTCTAGTCGAAA |
| p35S TATA | TGAGACTTTTCAACAAAGGGTAATATCGGGAAACCTCCTCGGATTCCATTGCCCAGCTATCTGTCACTTCATCAAAAGGACAGTAGAAAAGGAAGGTGGCACCTACAAATGCCATCATTGCGATAAAGGAAAGGCTATCGTTCAAGATGCCCCTGCCGACAGTGGTCCCAAAGATGGACCCCCACCCACGAGGAGCATCGTGGAAAAAGAAGACGTTCCAACCACGTCTTCAAAGCAAGTGGATTGATGTGATATCTCCACTGACGTAAGGGATGACGCACAATCCCACTATCCTTCGCAAGACCCTTCCTC**AATTATATACTACTATTAGTACCTAGTCTTAATT**TATATAAGGAAGTTCATTTCATTTGGAGAGGACTCCGGTATTTTTACAACAATTACCACAACAAAACAAACAACAAACAACATTACAATTTACTATTCTAGTCGAAA |
| p35S 3’ | TGAGACTTTTCAACAAAGGGTAATATCGGGAAACCTCCTCGGATTCCATTGCCCAGCTATCTGTCACTTCATCAAAAGGACAGTAGAAAAGGAAGGTGGCACCTACAAATGCCATCATTGCGATAAAGGAAAGGCTATCGTTCAAGATGCCCCTGCCGACAGTGGTCCCAAAGATGGACCCCCACCCACGAGGAGCATCGTGGAAAAAGAAGACGTTCCAACCACGTCTTCAAAGCAAGTGGATTGATGTGATATCTCCACTGACGTAAGGGATGACGCACAATCCCACTATCCTTCGCAAGACCCTTCCTCTATATAAGGAAGTTCATTTCATTTGGAGAGGACTCCGGTATTTTTACAACAATTACCACAACAAAACAAACAACAAACAACATTACAATTTACTATTCTAGTCGAA**AATTATATACTACTATTAGTACCTAGTCTTAATT**A |
| p35S TATA+3’ | TGAGACTTTTCAACAAAGGGTAATATCGGGAAACCTCCTCGGATTCCATTGCCCAGCTATCTGTCACTTCATCAAAAGGACAGTAGAAAAGGAAGGTGGCACCTACAAATGCCATCATTGCGATAAAGGAAAGGCTATCGTTCAAGATGCCCCTGCCGACAGTGGTCCCAAAGATGGACCCCCACCCACGAGGAGCATCGTGGAAAAAGAAGACGTTCCAACCACGTCTTCAAAGCAAGTGGATTGATGTGATATCTCCACTGACGTAAGGGATGACGCACAATCCCACTATCCTTCGCAAGACCCTTCCTC**AATTATATACTACTATTAGTACCTAGTCTTAATT**TATATAAGGAAGTTCATTTCATTTGGAGAGGACTCCGGTATTTTTACAACAATTACCACAACAAAACAAACAACAAACAACATTACAATTTACTATTCTAGTCGAA**TAGAATTAGTACCTGATACTAATAATT**A |
| p35S 2xTATA | TGAGACTTTTCAACAAAGGGTAATATCGGGAAACCTCCTCGGATTCCATTGCCCAGCTATCTGTCACTTCATCAAAAGGACAGTAGAAAAGGAAGGTGGCACCTACAAATGCCATCATTGCGATAAAGGAAAGGCTATCGTTCAAGATGCCCCTGCCGACAGTGGTCCCAAAGATGGACCCCCACCCACGAGGAGCATCGTGGAAAAAGAAGACGTTCCAACCACGTCTTCAAAGCAAGTGGATTGATGTGATATCTCCACTGACGTAAGGGATGACGCACAATCCCACTATCCTTCGCAAGACCCTTCCTC**TAGAATTAGTACCTGATACTAATAATT**GAGGCACGAC**AATTATATACTACTATTAGTACCTAGTCTTAATT**TATATAAGGAAGTTCATTTCATTTGGAGAGGACTCCGGTATTTTTACAACAATTACCACAACAAAACAAACAACAAACAACATTACAATTTACTATTCTAGTCGAAA |
| NahR | Meyer *et al.,* 2019 |

| | |
|---|---|
| p35S 5’ | **CGCAATATTCATGTTGATGATTTATTATATA**TGAGACTTTTCAACAAAGGGTAATATCGGGAAACCTCCTCGGATTCCATTGCCCAGCTATCTGTCACTTCATCAAAAGGACAGTAGAAAAGGAAGGTGGCACCTACAAATGCCATCATTGCGATAAAGGAAAGGCTATCGTTCAAGATGCCCCTGCCGACAGTGGTCCCAAAGATGGACCCCCACCCACGAGGAGCATCGTGGAAAAAGAAGACGTTCCAACCACGTCTTCAAAGCAAGTGGATTGATGTGATATCTCCACTGACGTAAGGGATGACGCACAATCCCACTATCCTTCGCAAGACCCTTCCTCTATATAAGGAAGTTCATTTCATTTGGAGAGGACTCCGGTATTTTTACAACAATTACCACAACAAAACAAACAACAAACAACATTACAATTTACTATTCTAGTCGAAA |
| p35S TATA | TGAGACTTTTCAACAAAGGGTAATATCGGGAAACCTCCTCGGATTCCATTGCCCAGCTATCTGTCACTTCATCAAAAGGACAGTAGAAAAGGAAGGTGGCACCTACAAATGCCATCATTGCGATAAAGGAAAGGCTATCGTTCAAGATGCCCCTGCCGACAGTGGTCCCAAAGATGGACCCCCACCCACGAGGAGCATCGTGGAAAAAGAAGACGTTCCAACCACGTCTTCAAAGCAAGTGGATTGATGTGATATCTCCACTGACGTAAGGGATGACGCACAATCCCACTATCCTTCGCAAGACCCTTCCTC**CGCAATATTCATGTTGATGATTTATTATATA**TATATAAGGAAGTTCATTTCATTTGGAGAGGACTCCGGTATTTTTACAACAATTACCACAACAAAACAAACAACAAACAACATTACAATTTACTATTCTAGTCGAAA |
| p35S 3’ | TGAGACTTTTCAACAAAGGGTAATATCGGGAAACCTCCTCGGATTCCATTGCCCAGCTATCTGTCACTTCATCAAAAGGACAGTAGAAAAGGAAGGTGGCACCTACAAATGCCATCATTGCGATAAAGGAAAGGCTATCGTTCAAGATGCCCCTGCCGACAGTGGTCCCAAAGATGGACCCCCACCCACGAGGAGCATCGTGGAAAAAGAAGACGTTCCAACCACGTCTTCAAAGCAAGTGGATTGATGTGATATCTCCACTGACGTAAGGGATGACGCACAATCCCACTATCCTTCGCAAGACCCTTCCTCTATATAAGGAAGTTCATTTCATTTGGAGAGGACTCCGGTATTTTTACAACAATTACCACAACAAAACAAACAACAAACAACATTACAATTTACTATTCTAGTCGAA**CGCAATATTCATGTTGATGATTTATTATATA**A |
| p35S 2xTATA | TGAGACTTTTCAACAAAGGGTAATATCGGGAAACCTCCTCGGATTCCATTGCCCAGCTATCTGTCACTTCATCAAAAGGACAGTAGAAAAGGAAGGTGGCACCTACAAATGCCATCATTGCGATAAAGGAAAGGCTATCGTTCAAGATGCCCCTGCCGACAGTGGTCCCAAAGATGGACCCCCACCCACGAGGAGCATCGTGGAAAAAGAAGACGTTCCAACCACGTCTTCAAAGCAAGTGGATTGATGTGATATCTCCACTGACGTAAGGGATGACGCACAATCCCACTATCCTTCGCAAGACCCTTCCTC**CGCAGTATTCATGCTGATGATTTATTATATA**GTCGTCGGTC**CGCAATATTCATGTTGATGATTTATTATATA**TATATAAGGAAGTTCATTTCATTTGGAGAGGACTCCGGTATTTTTACAACAATTACCACAACAAAACAAACAACAAACAACATTACAATTTACTATTCTAGTCGAAA |
| p35S 3x5’ | **CGCAGTATTCACGCTGGTGATAAACAATTC**CAGTTCCTTA**CGCAGTATTCATGCTGATGATTTATTATATA**CTGTGCACTA**CGCAATATTCATGTTGATGATTTATTATATA**TGAGACTTTTCAACAAAGGGTAATATCGGGAAACCTCCTCGGATTCCATTGCCCAGCTATCTGTCACTTCATCAAAAGGACAGTAGAAAAGGAAGGTGGCACCTACAAATGCCATC |

| | |
|---|---|
| | ATTGCGATAAAGGAAAGGCTATCGTTCAAGATGCCCCTGCCGACAGTGGTCCCAAAGATGGACCCCCACCCACGAGGAGCATCGTGGAAAAAGAAGACGTTCCAACCACGTCTTCAAAGCAAGTGGATTGATGTGATATCTCCACTGACGTAAGGGATGACGCACAATCCCACTATCCTTCGCAAGACCCTTCCTCTATATAAGGAAGTTCATTTCATTTGGAGAGGACTCCGGTATTTTTACAACAATTACCACAACAAAACAAACAACAAACAACATTACAATTTACTATTCTAGTCGAAA |
| p35S 4x | **CGCAGTATTCACGCTGGTGATAAACAAATTC**CAGTTCCTTA**CGCAGTATTCATGCTGATGATTTATTATATA**CTGTGCACTA**CGCAATATTCATGTTGATGATTTATTATATA**TGAGACTTTTCAACAAAGGGTAATATCGGGAAACCTCCTCGGATTCCATTGCCCAGCTATCTGTCACTTCATCAAAAGGACAGTAGAAAAGGAAGGTGGCACCTACAAATGCCATCATTGCGATAAAGGAAAGGCTATCGTTCAAGATGCCCCTGCCGACAGTGGTCCCAAAGATGGACCCCCACCCACGAGGAGCATCGTGGAAAAAGAAGACGTTCCAACCACGTCTTCAAAGCAAGTGGATTGATGTGATATCTCCACTGACGTAAGGGATGACGCACAATCCCACTATCCTTCGCAAGACCCTTCCTC**CGCAATATTCATGTTGATGATTTATTATATA**TATATAAGGAAGTTCATTTCATTTGGAGAGGACTCCGGTATTTTTACAACAATTACCACAACAAAACAAACAACAAACAACATTACAATTTACTATTCTAGTCGAAA |
| p35S 5x | **CGCAGTATTCACGCTGGTGATAAACAAATTC**CAGTTCCTTA**CGCAGTATTCATGCTGATGATTTATTATATA**CTGTGCACTA**CGCAATATTCATGTTGATGATTTATTATATA**TGAGACTTTTCAACAAAGGGTAATATCGGGAAACCTCCTCGGATTCCATTGCCCAGCTATCTGTCACTTCATCAAAAGGACAGTAGAAAAGGAAGGTGGCACCTACAAATGCCATCATTGCGATAAAGGAAAGGCTATCGTTCAAGATGCCCCTGCCGACAGTGGTCCCAAAGATGGACCCCCACCCACGAGGAGCATCGTGGAAAAAGAAGACGTTCCAACCACGTCTTCAAAGCAAGTGGATTGATGTGATATCTCCACTGACGTAAGGGATGACGCACAATCCCACTATCCTTCGCAAGACCCTTCCTC**CGCAGTATTCATGCTGATGATTTATTATATA**GTCGTCGGTC**CGCAATATTCATGTTGATGATTTATTATATA**TATATAAGGAAGTTCATTTCATTTGGAGAGGACTCCGGTATTTTTACAACAATTACCACAACAAAACAAACAACAAACAACATTACAATTTACTATTCTAGTCGAAA |
| PadR | Nguyen *et al.,* 2011; Park *et al.,* 2017 |
| p35S TATA | TGAGACTTTTCAACAAAGGGTAATATCGGGAAACCTCCTCGGATTCCATTGCCCAGCTATCTGTCACTTCATCAAAAGGACAGTAGAAAAGGAAGGTGGCACCTACAAATGCCATCATTGCGATAAAGGAAAGGCTATCGTTCAAGATGCCCCTGCCGACAGTGGTCCCAAAGATGGACCCCCACCCACGAGGAGCATCGTGGAAAAAGAAGACGTTCCAACCACGTCTTCAAAGCAAGTGGATTGATGTGATATCTCCACTGACGTAAGGGATGACGCACAATCCCACTATCCTTCGCAAGACCCTTCCTC**AACATGTAAATAGTTACATGAT**TATATAAGGAAGTTCATTTCATTTGGAGAGGACTCCGGTATTTTTACAACAATTACCACAACAAAACAAACAACAAACAACATTACAATTTACTATTCTAGTCGAAA |

| | |
|---|---|
| p35S 2xTATA | TGAGACTTTTCAACAAAGGGTAATATCGGGAAACCTCCTCGGATTCC<br>ATTGCCCAGCTATCTGTCACTTCATCAAAAGGACAGTAGAAAAGGA<br>AGGTGGCACCTACAAATGCCATCATTGCGATAAAGGAAAGGCTATC<br>GTTCAAGATGCCCCTGCCGACAGTGGTCCCAAAGATGGACCCCCAC<br>CCACGAGGAGCATCGTGGAAAAAGAAGACGTTCCAACCACGTCTTC<br>AAAGCAAGTGGATTGATGTGATATCTCCACTGACGTAAGGGATGAC<br>GCACAATCCCACTATCCTTCGCAAGACCCTTCCTC**ATGGTGTTAAA**<br>**GTGAACATGTAAATAGTTACATGAT**TATATAAGGAAGTTCATTTCA<br>TTTGGAGAGGACTCCGGTATTTTTACAACAATTACCACAACAAAACA<br>AACAACAAACAACATTACAATTTACTATTCTAGTCGAAA |
| p35S TATA+3’ | TGAGACTTTTCAACAAAGGGTAATATCGGGAAACCTCCTCGGATTCC<br>ATTGCCCAGCTATCTGTCACTTCATCAAAAGGACAGTAGAAAAGGA<br>AGGTGGCACCTACAAATGCCATCATTGCGATAAAGGAAAGGCTATC<br>GTTCAAGATGCCCCTGCCGACAGTGGTCCCAAAGATGGACCCCCAC<br>CCACGAGGAGCATCGTGGAAAAAGAAGACGTTCCAACCACGTCTTC<br>AAAGCAAGTGGATTGATGTGATATCTCCACTGACGTAAGGGATGAC<br>GCACAATCCCACTATCCTTCGCAAGACCCTTCCTC**AACATGTAAATA**<br>**GTTACATGAT**TATATAAGGAAGTTCATTTCATTTGGAGAGGACTCC<br>GGTATTTTTACAACAATTACCACAACAAAACAAACAACAAACAACA<br>TTACAATTTACTATTCTAGTCGAA**ATGGTGTTAAAGTGAACATGTA**<br>A |
| TtgR | Meyer *et al.,* 2019 |
| p35S 5’ | **TATTTACAAACAACCATGAATGTAAGTA**TGAGACTTTTCAACAAA<br>GGGTAATATCGGGAAACCTCCTCGGATTCCATTGCCCAGCTATCTGT<br>CACTTCATCAAAAGGACAGTAGAAAAGGAAGGTGGCACCTACAAAT<br>GCCATCATTGCGATAAAGGAAAGGCTATCGTTCAAGATGCCCCTGC<br>CGACAGTGGTCCCAAAGATGGACCCCCACCCACGAGGAGCATCGTG<br>GAAAAAGAAGACGTTCCAACCACGTCTTCAAAGCAAGTGGATTGAT<br>GTGATATCTCCACTGACGTAAGGGATGACGCACAATCCCACTATCCT<br>TCGCAAGACCCTTCCTCTATATAAGGAAGTTCATTTCATTTGGAGAG<br>GACTCCGGTATTTTTACAACAATTACCACAACAAAACAAACAACAA<br>ACAACATTACAATTTACTATTCTAGTCGAAA |
| p35S TATA | TGAGACTTTTCAACAAAGGGTAATATCGGGAAACCTCCTCGGATTCC<br>ATTGCCCAGCTATCTGTCACTTCATCAAAAGGACAGTAGAAAAGGA<br>AGGTGGCACCTACAAATGCCATCATTGCGATAAAGGAAAGGCTATC<br>GTTCAAGATGCCCCTGCCGACAGTGGTCCCAAAGATGGACCCCCAC<br>CCACGAGGAGCATCGTGGAAAAAGAAGACGTTCCAACCACGTCTTC<br>AAAGCAAGTGGATTGATGTGATATCTCCACTGACGTAAGGGATGAC<br>GCACAATCCCACTATCCTTCGCAAGACCCTTCCTC**TATTTACAAACA**<br>**ACCATGAATGTAAGTA**TATATAAGGAAGTTCATTTCATTTGGAGAG<br>GACTCCGGTATTTTTACAACAATTACCACAACAAAACAAACAACAA<br>ACAACATTACAATTTACTATTCTAGTCGAAA |
| p35S 3’ | TGAGACTTTTCAACAAAGGGTAATATCGGGAAACCTCCTCGGATTCC<br>ATTGCCCAGCTATCTGTCACTTCATCAAAAGGACAGTAGAAAAGGA<br>AGGTGGCACCTACAAATGCCATCATTGCGATAAAGGAAAGGCTATC<br>GTTCAAGATGCCCCTGCCGACAGTGGTCCCAAAGATGGACCCCCAC |

| | |
|---|---|
| | CCACGAGGAGCATCGTGGAAAAAGAAGACGTTCCAACCACGTCTTC<br>AAAGCAAGTGGATTGATGTGATATCTCCACTGACGTAAGGGATGAC<br>GCACAATCCCACTATCCTTCGCAAGACCCTTCCTCTATATAAGGAAG<br>TTCATTTCATTTGGAGAGGACTCCGGTATTTTTACAACAATTACCAC<br>AACAAAACAAACAACAAACAACATTACAATTTACTATTCTAGTCGA<br>A**TATTTACAAACAACCATGAATGTAAGTA**A |
| p35S 2xTATA | TGAGACTTTTCAACAAAGGGTAATATCGGGAAACCTCCTCGGATTCC<br>ATTGCCCAGCTATCTGTCACTTCATCAAAAGGACAGTAGAAAAGGA<br>AGGTGGCACCTACAAATGCCATCATTGCGATAAAGGAAAGGCTATC<br>GTTCAAGATGCCCCTGCCGACAGTGGTCCCAAAGATGGACCCCCAC<br>CCACGAGGAGCATCGTGGAAAAAGAAGACGTTCCAACCACGTCTTC<br>AAAGCAAGTGGATTGATGTGATATCTCCACTGACGTAAGGGATGAC<br>GCACAATCCCACTATCCTTCGCAAGACCCTTCCTC**TATATACATACA**<br>**TGCATGTATGTATGTA**GAGGCACGAC**TATTTACAAACAACCATGA**<br>**ATGTAAGTA**TATATAAGGAAGTTCATTTCATTTGGAGAGGACTCCG<br>GTATTTTTACAACAATTACCACAACAAAACAAACAACAAACAACAT<br>TACAATTTACTATTCTAGTCGAAA |
| p35S TATA+3’ | TGAGACTTTTCAACAAAGGGTAATATCGGGAAACCTCCTCGGATTCC<br>ATTGCCCAGCTATCTGTCACTTCATCAAAAGGACAGTAGAAAAGGA<br>AGGTGGCACCTACAAATGCCATCATTGCGATAAAGGAAAGGCTATC<br>GTTCAAGATGCCCCTGCCGACAGTGGTCCCAAAGATGGACCCCCAC<br>CCACGAGGAGCATCGTGGAAAAAGAAGACGTTCCAACCACGTCTTC<br>AAAGCAAGTGGATTGATGTGATATCTCCACTGACGTAAGGGATGAC<br>GCACAATCCCACTATCCTTCGCAAGACCCTTCCTC**TATTTACAAACA**<br>**ACCATGAATGTAAGTA**TATATAAGGAAGTTCATTTCATTTGGAGAG<br>GACTCCGGTATTTTTACAACAATTACCACAACAAAACAAACAACAA<br>ACAACATTACAATTTACTATTCTAGTCGAA**TATATACATACATGCA**<br>**TGTATGTATGTA**A |

**Table S3** Primers used in this study.

| Name | Sequence | Notes |
|---|---|---|
| ACC1 penny Fw1 | ATTACGTCTCGCTCGAATGGCTGGCTCCATTAACGGG | ACC1 cloning |
| ACC1 penny Rv1 | ATTACGTCTCACTCACCGAAACCACGGCAGCACTGGTCG | ACC1 cloning |
| ACC1 penny Fw2 | TGTGCGTCTCTCTCGTCGGACTACGTTGGTTATCTTGAGAAGG | ACC1 cloning |
| ACC1 penny Rv2 | ATCTCGTCTCGCTCAGCCACCATTCCAAATTCATTAAGACCCG | ACC1 cloning |
| ACC1 penny Fw3 | ATTACGTCTCACTCGTGGCCTGGTGCCTTGAGATG | ACC1 cloning |
| ACC1 penny Rv3 | TAGCCGTCTCGCTCAAAGCTCAACCCAACACCTTTCGTAGCG | ACC1 cloning |

| | | |
|---|---|---|
| AAE13 FW | AGTCCGTCTCGCTCGAATGACCGCTACGACAACATTAAAG | AAE13 cloning |
| AAE13 RV | ATTCCGTCTCGCTCAAAGCTCACTCTTGATTTTCCAGAGATTTCTTTAG | AAE13 cloning |
| AAE13 SV FW | AGTCCGTCTCGCTCGAATGGAAGTGTTTAAAGCAGCTT | AAE13 SV cloning |

**Table S4** Nucleotide sequences of additional domains fused to aTFs, promoters, and other protein fusions not built in this work. Sequences were retrieved from the references or accession number as indicated below.

| | |
|---|---|
| SRDX | Golden braid 2.0 Kit Sarrion-Perdigones *et al.,* 2013 |
| TTGGACCTTGATCTTGAATTGAGACTTGGTTTTGCA | |
| NLS | Golden braid 2.0 Kit Sarrion-Perdigones *et al.,* 2013 |
| CCAAAAAAGAAGCGTAAGGTT | |
| pGal4-min35S (Gal4 UAS in bold, p35S -46 to +1 underlined) | Golden braid 2.0 Kit Sarrion-Perdigones *et al.,* 2013 |
| **CGGAGTACTGTCCTCCG**AG**CGGAGTACTGTCCTCCG**AG**CGGAGTACTGTCCTCCG**AG**CGGAGTACTGTCCTCCG**AG**CGGAGTACTGTCCTCCG**TCCCCTTCGCAAGACCCTTCCTCTATATAAGGAAGTTCATTTCATTTGGAGAGGACTCCGGTATTTTTACAACAATTACCACAACAAAACAAACAACAAACAACATTACAATTTACTATTCTAGTCGAA | |
| PEST Sequence (flexible linker underlined) | pNL1.2[NlucP] vector, Genebank JQ437371.1 |
| TCGGGGTCCGGCAGCCACGGTTTTCCACCTGAGGTCGAGGAACAGGCGGCAGGAACCCTGCCCATGTCCTGCGCTCAGGAGTCTGGTATGGACAGACATCCCGCTGCATGTGCAAGCGCCAGAATTAACGTG | |
| NLS-**Gal4**-linker-**VP64** fusion | Anderson *et al.,* 2023 |
| ATGCCTAAAAAAAAACGTAAGGTA**ATGAAGCTACTGTCTTCTATCGAACAAGCATGCGATATTTGCCGACTTAAAAAGCTCAAGTGCTCCAAAGAAAAACCGAAGTGCGCCAAGTGTCTGAAGAACAACTGGGAGTGTCGCTACTCTCCCAAAACCAAAAGATCTCCGCTGACTAGGGCACATCTGACAGAAGTGGAATCAAGGCTAGAAAGACTGGAACAGCTATTTCTACTGATTTTTCCTAGG**GGCGGCAGCGGAGGCTCTGGTTCG**GACGCATTGGATGACTTCGACTTAGACATGCTTGGGTCAGATGCTTTGGACGACTTTGACCTCGACATGTTGGGTTCTGACGCGCTAGATGACTTCGACTTGGACATGTTGGGCTCCGACGCTTTGGACGACTTTGATCTGGACATGTTATAA** | |
| **Gal4**-NLS-P4 fusion | Anderson *et al.,* 2023 |
| **ATGAAGCTACTGTCTTCTATCGAACAAGCATGCGATATTTGCCGACTTAAAAAGCTCAAGTGCTCCAAAGAAAAACCGAAGTGCGCCAAGTGTCTGAAGAACAACTGGGAGTGTCGCTACTCTCCCAAAACCAAAAGATCTCCGCTGACTAGGGCACATCTGACAGAAGTGGAATCAAGGCTAGAAAGACTGGAACAGCTATTTCTACTGATTTTTCCTAGG**CCTAAAAAAAAACGTAAGGTAGGTTCGGGGGGTTCCGGTGG | |

| | |
|---|---|
| TTCTTCCCCTGAAGATAAGATTGCACAGCTCAAGCAGAAAATCCAAGCGTTAAAA<br>CAGGAGAACCAACAGCTAGAGGAAGAGAATGCTGCGTTGGAATATGGTTAA | |
| P3-NLS-**VP64** fusion | Anderson *et al.,* 2023 |
| ATGAGCCCGGAAGATGAGATCCAGCAACTGGAAGAAGAGATCGCCCAGCTAGAG<br>CAGAAGAACGCTGCGTTAAAAGAAAAGAATCAGGCACTAAAATATGGCGGGGT<br>TCCGGTGGTTCTTCAATGCCTAAAAAAAACGTAAGGTA**GACGCATTGGATGAC**<br>**TTCGACTTAGACATGCTTGGGTCAGATGCTTTGGACGACTTTGACCTCGACA**<br>**TGTTGGGTTCTGACGCGCTAGATGACTTCGACTTGGACATGTTGGGCTCCGA**<br>**CGCTTTGGACGACTTTGATCTGGACATGTTATAA** | |

**Table S5** Promoter constructs that showed the highest repression level for each respective aTF.

| aTF | Promoter construct | Repression level |
|---|---|---|
| CouR | p35S 5x | 95% (with SRDX-CouR fusion) |
| FapR | p35S TATA | 95% |
| PadR | p35S TATA | 23% |
| NahR | p35S 5x | 58% |
| TtgR | p35S 2xTATA | 96% |
